# Analysis of *Aedes aegypti* microRNAs in response to *Wolbachia w*AlbB infection and their potential role in mosquito longevity

**DOI:** 10.1101/2022.04.26.488845

**Authors:** Cameron Bishop, Mazhar Hussain, Leon E. Hugo, Sassan Asgari

## Abstract

The mosquito *Aedes aegypti* is the primary vector of a range of medically important viruses including dengue, Zika, West Nile, yellow fever, and chikungunya viruses. The endosymbiotic bacterium *Wolbachia pipientis w*AlbB strain is a promising biocontrol agent for blocking viral transmission by *Ae. aegypti*. To predict the long-term efficacy of field applications, a thorough understanding of the interactions between symbiont, host, and pathogen is required. *Wolbachia* influence host physiology in a variety of ways including reproduction, immunity, metabolism, and longevity. MicroRNAs (miRNAs) are highly conserved small non-coding RNAs that regulate gene expression in eukaryotes and viruses. A number of miRNAs are known to regulate biological processes in *Drosophila* and mosquitoes, including facilitating *Wolbachia* maintenance. We generated the first chromosomal map of *Ae. aegypti* miRNAs, and compared miRNA expression profiles between a *w*AlbB-transinfected *Ae. aegypti* mosquito line and a tetracycline cleared derivative, using deep small RNA-sequencing. We found limited modulation of miRNAs in response to *w*AlbB infection. Several miRNAs were modulated in response to age, some of which showed greater upregulation in *w*AlbB-infected mosquitos than in tetracycline cleared ones. By selectively inhibiting some differentially expressed miRNAs, we identified miR-2946-3p and miR-317-3p as effecting mosquito longevity.

**Importance:** *Wolbachia* is an endosymbiotic bacterium found in about 65% of insect species. It is mostly known for reproductive manipulations of the host, and also blocking replication of positive sense RNA viruses. Transinfection of *Wolbachia* into *Aedes aegypti* mosquitoes, which transmit a variety of arboviruses, including dengue virus, has provided a novel biological approach in reducing transmission of arboviruses. To gain a better understanding of *Wolbachia*-mosquito interactions, we investigated the impact of *Wolbachia* on the microRNA profile of *Ae. aegypti* mosquitoes. We produced the first chromosome-level map of *Ae. Aegypti* miRNAs. We found modulation of microRNAs in mosquitoes due to age, with two miRNAs, 317-3p and 2946-3p, showing significant increase with age. Inhibition of 317-3p and 2946-3p led to reduced mosquito life span in *w*AlbB-infected mosquitoes. The outcomes provide insights into underlying molecular mechanisms involved in *Wolbachia*-host interactions.

## Introduction

Arthropod-borne viruses (arboviruses) pose a considerable and increasing global health burden. Two mosquitoes of the *Aedes* genus, *Aedes aegypti* and *Aedes albopictus,* are the primary and secondary vectors, respectively, of several medically important arboviruses including the flaviviruses dengue virus (DENV), Zika virus (ZIKV), yellow fever virus (YFV), and the alphavirus chikungunya virus (CHIKV) ^1, 2^. Originating in sub-Saharan Africa, *Ae. aegypti* experienced a dramatic range expansion beginning with European colonisation of the Americas in the 16^th^ century ^3^. Human activity has continued to facilitate its global spread, and today it is common in many urban subtropical and tropical habitats across six continents ^4^. In the absence of safe or effective vaccines, vector control remains the most effective method for limiting outbreaks of mosquito-borne viruses ^5, 6^. With many *Ae. aegypti* populations becoming resistant to insecticides, attention has turned to biological vector-control strategies ^7–9^.

The endosymbiotic bacterium *Wolbachia pipientis* has proven to be a useful biocontrol agent through its ability to invade mosquito populations and prevent viral transmission, either by suppressing the mosquito population or by inhibiting virus infection and dissemination within individual mosquitoes ^10, 11^. The ability for some strains of *Wolbachia* to invade mosquito populations is due to a combination of maternal inheritance and a manipulation by the bacterium known as cytoplasmic incompatibility, which provides a reproductive advantage to infected females in naïve or heterologously-infected populations ^12, 13^. Additional factors governing the success of a *Wolbachia* invasion include the fitness cost to the mosquito harbouring *Wolbachia*, and the ability of *Wolbachia* to tolerate the climate of tropical regions, where mosquito-borne viruses primarily exist ^14–16^. The viral-inhibition phenotype exhibited by some strains of *Wolbachia* is complex, not well-understood, and varies depending on the combination of *Wolbachia* strain and mosquito host ^17–19^.

Several *Wolbachia* strains have been investigated for their efficacy in biocontrol, and to explore the biology that underpins their association with the host. The *Drosophila*-derived *w*MelPop and *w*Mel strains have been investigated in laboratory and/or field trials for suppression of West Nile virus (WNV), YFV, ZIKV, DENV, and CHIKV in *Ae. aegypti*. Despite effectively supressing viral infection and transmission ^20–24^, their ability to be maintained at high frequency in wild populations is limited by the large fitness cost of *w*MelPop ^14^, and the sensitivity to heat-stress of *w*Mel ^25, 26^. An *Ae. albopictus*-derived strain, *w*AlbB, inhibits transmission of DENV in *Ae. aegypti*, while also being more tolerant of heat stress than *w*Mel or *w*MelPop, allowing it to persist in wild populations ^25, 27–30^.

The symbiotic relationship between *Wolbachia* and host is intimate and complex, with each party affecting the other through multiple physiological pathways, including innate immunity, redox homeostasis, metabolism, protein synthesis and proteolysis, nutrient provisioning, and iron homeostasis ^19, 31–39^. *Wolbachia* infection has also been shown to have a significant effect on host gene expression via the microRNA (miRNA) pathway.

miRNAs are small non-coding RNAs involved in the regulation of gene expression in eukaryotes and viruses ^40^. In mosquitoes, a number of miRNAs are conserved among disparate lineages, indicating their importance in regulating critical functions. For example, miR-281, miR-184, miR-989 and miR-278 are conserved among *Anopheles gambiae*, *Ae. aegypti*, and *Culex quinquefasciatus* ^41^. Some conserved miRNAs also show conserved patterns of expression among tissue types, or developmental stages, or in response to physiological cues such as blood feeding, reproduction, or infection by pathogens ^41, 42^. In *Ae. aegypti*, several miRNAs have been functionally explored. The ovary-specific miR-309 is induced by blood feeding and has been shown to regulate ovarian development by targeting the *SIX4* transcript ^43^. miR-1174 is involved in bloodmeal intake by targeting *serine hydroxymethyltransferase*, while miR-1890 regulates blood digestion by targeting the *JHA15* transcript ^41, 44^.

A number of miRNAs are modulated by arbovirus infection in *Ae. aegypti*, some of which have been predicted to have an effect on viral replication ^41, 45^. For example, miR-2944b-5p is exploited by CHIKV to enhance its replication via a direct interaction with the 3’ UTR of the viral genome, and indirectly via repression of a host target gene, vps-13 ^46^. Similarly, the blood-meal induced miR-375 has been implicated in enhancing DENV-2 infection in *Ae. aegypti* via the Toll pathway by inducing *cactus* and repressing *REL1* ^47^. In experiments using *Ae. aegypti*, *Wolbachia w*MelPop induced the expression of miR-2940, which in turn promoted the maintenance of *w*MelPop by supressing methyltransferase *Dnmt2*, and by inducing arginine methyltransferase ^48, 49^. Furthermore, miR-2940 has been shown to inhibit DENV and WNV infection in *Ae. aegypti* and *Ae. albopictus* via induction of the metalloprotease m41 ftsh ^48, 50^. The miR-2940 instance provides an elegant example of symbiosis between *Wolbachia* and *Ae. aegypti,* and underscores the importance of the miRNA pathway in facilitating the interaction. It remains to be seen how the effect on host miRNA profile imposed by *w*MelPop compares to that of *w*AlbB.

To further shed light on the role of miRNAs in mosquito-*Wolbachia* interactions, we used deep-sequencing of small RNAs to investigate the effect of *Wolbachia w*AlbB strain on the miRNA expression profile of *Ae. aegypti* mosquitoes. By comparing *w*AlbB-infected mosquitoes with a tetracycline-cleared line, we detected limited differential expression of miRNAs in response to *w*AlbB infection. The effect of age was stronger than that of *w*AlbB infection, and some miRNAs were upregulated with age in a manner that appeared to be exaggerated by *w*AlbB. Further, our results suggest that upregulation of aae-miR-2946-3p and aae-miR-317-3p could be related to mosquito longevity in *w*AlbB-infected mosquitoes. A number of the differentially expressed miRNAs have been described elsewhere as being involved in the regulation of metabolism, microbial defence, reproduction, development and ageing in mosquitoes.

## Materials and Methods

### Mosquitoes

The *Wolbachia w*AlbB infected *Ae. aegypti* strain *w*AlbB2-F4 mosquitoes ^51^ were used for experiments in this study. We call the strain WB2 for shortness. A *Wolbachia*-free (WB2.tet) line was produced by feeding adults on 10% sucrose with 1mg/ml tetracycline hydrochloride (Sigma) for seven generations. WB2.tet mosquitoes were maintained for five generations without tetracycline before testing for the presence of *Wolbachia w*AlbB using qPCR on genomic DNA with primers to the *Wolbachia surface protein* (*wsp*) gene from nine adult females to ensure removal of *Wolbachia*. See qPCR below for primer sequences and details. Female WB2 mosquitoes were also tested for *Wolbachia* density by the same method. Both lines were reared by hatching 300 eggs in 27°C water and feeding larvae daily with ground Tropical Colour flasks (Tetra) fish food *ad libitum.* Adults were maintained at 27°C, a 12hr:12hr day/night cycle with relative humidity ranging between 65-75%, and allowed to feed *ad libitum* on 10% sucrose. All mosquito experiments were performed using female adults.

### Cell line

The *Ae. aegypti* Aag2.*w*AlbB cell line was generated as described in ^52^ for testing the effect of miRNA inhibition of aae-miR-2b-3p, aae-miR-190-5p, and aae-miR-276b-5p on *w*AlbB density. Cells were maintained in a 1:1 ratio of Mitsuhashi-Maramorosch (Himedia) and Schneider’s Drosophila Medium (Invitrogen, Carlsbad, USA) supplemented with 10% Foetal Bovine Serum (FBS) (Bovogen Biologicals, French origin) at 27°C as monolayer.

### Small RNA sequencing

Adult female mosquitoes were sampled at 2, 6, and 12 days post emergence. Three replicate pools of four females were sampled at each time point. Samples were homogenised in 1.5ml tubes on ice using a plastic micro-pestle and small RNAs were extracted using the miRNeasy Mini Kit (QIAGEN, Hilden, Germany). Genomic DNA was removed using the Turbo DNA-free kit (Invitrogen) as per the manufacturer’s instructions. Total RNA from each sample was quantified and purity determined using an Agilent 2100 Bioanalyzer (Agilent Technologies, Palo Alto, CA, USA), NanoDrop (Thermo Fisher Scientific Inc.) and performing gel electrophoresis in a 1% agarose gel. The following steps relating to library preparation and sequencing were carried out by the sequencing service company Genewiz (China). Two µg of total RNA with RNA integrity number (RIN) value of above 7.5 was used for subsequent library preparation. Indexed sequencing libraries were constructed according to the manufacturer’s protocol (NEBNext Multiplex Small RNA library Prep Set for Illumina). The resulting PCR products of ∼140 bp were recovered and purified via polyacrylamide agarose gel electrophoresis, validated using an Agilent 2100 Bioanalyzer (Agilent Technologies), and quantified using a Qubit 2.0 Fluorometer (Invitrogen). Libraries were then multiplexed and sequenced on an Illumina HiSeq 2500 instrument using a 1x50 single-end configuration according to manufacturer’s instructions (Illumina, San Diego, CA, USA). Image analysis and base calling were conducted using the HiSeq Control Software (HCS), Off-Line Basecaller (OLB), and GAPipeline-1.6 (Illumina).

### Bioinformatics analyses

Low quality reads and adapters were removed from raw small RNA sequence reads using Trimmomatic. Cleaned reads were then size-selected for reads between 18-24nt using BBmap ^53^. Quality control was performed using FastQC ^54^. Mature miRNA annotations were generated with miRdeep2 v0.1.2 ^55^ using four concatenated read files (two WB2.tet and two WB2 - SRR12893564, SRR12893577, SRR12893581, SRR12893570), a previously published list of *Ae. aegypti* precursor miRNA sequences ^42^, and miRBase databases for *Ae. aegypti*, *Anopheles gambiae*, and *Drosophila melanogaster* ^56^ as input. miRDeep2 output provides genomic coordinates of precursor miRNA loci, but not their corresponding mature sequences, so we determined the exact mature miRNA locations using BLAST ^57^, and cross referenced the results with the precursor coordinates. This method also allowed -5p and -3p designation to each mature sequence, which was not included in miRDeep2’s output. To quantify miRNA expression, reads were mapped to the three complete chromosomes of the *Aedes aegypti* AaegL5.0 reference genome ^58^ using Shortstack v3.8.5 ^59^ with default settings. Reads mapping to mature miRNA annotations were counted using Shortstack with default settings. A differential expression analysis was performed using EdgeR v3.34.0 ^60^. Low-count miRNAs were removed using a minimum cutoff of 5 counts per million. Read counts were normalised for library depth and composition using the trimmed mean of M-values method ^61^. Genewise dispersion was modelled with the ‘robust=T’ parameter and the ‘glmFit’ function, and a test for differential expression was performed using the ‘glmLRT’ function. For contrasts between lines, miRNAs with a Benjamini & Hochberg adjusted *p* value of < 0.05 and a log_2_ fold-change > 0.5 were considered differentially expressed. For contrasts between ages, miRNAs with an adjusted *p* value of < 0.05 and a log_2_ fold-change of > 1 were considered differentially expressed. The decision to relax the log_2_ fold-change cut-off to 0.5 for contrasts between lines was based on the relatively low fold-changes observed in these contrasts compared to those in contrasts between ages. Positive log_2_ fold-changes indicate a higher expression level in the *w*AlbB-infected mosquitoes versus control. To assess the microbiome of each mosquito line, unmapped reads were analysed using Kraken with the minikraken2 database ^62^. All data were plotted using Kronatools ^63^ and R v3.6.3.

To predict gene targets of differentially expressed miRNAs, coding regions and 5’- and 3’- UTRs of the AaegL5.0 transcriptome were scanned for potential binding sites using three software packages: miRanda v3.3 ^64^, RNA22 v2.0 ^65^, and PITA v6.0 ^66^. We then took target genes predicted by all three packages and performed a gene ontology term enrichment analysis using VectorBase ^67^, with a *p* value cut-off of 0.05.

To identify putative hairpin structures expressed by *w*AlbB, WB2 fastq files were concatenated within each age group. Reads of between 18 and 24 nt in length were mapped to the *Wolbachia pipientis w*AlbB reference genome (Genbank Accession: NZ_CP031221.1) using ^68^ with bowtie2 v2.2.7 ^69^. Peaks of high coverage were identified (≥ 2000 reads) using bedtools v2.26.0 ^70^. High coverage peaks were visually examined using Integrated Genome Viewer v2.11.9. RNA secondary structure was predicted using the The ViennaRNA Package ^71^. Sequence alignment was performed using BioEdit v7.2 with the ‘sliding ends’ option.

### qPCR and RT-qPCR analysis

Genomic DNA was extracted from 3 pools of 4 adult females at 2, 6, and 12 days post-emergence using Econospin columns (Epoch Life Sciences), following a protocol described previously ^72^. DNA was quantified using a BioTek Epoch Microspot plate spectrophotometer. Quantitative PCR was performed using SYBR master mix (QIAGEN) in a QIAGEN Rotor-Gene Q 2plex using 100ng of gDNA as per the manufacturer’s instructions. The ratio of *Wolbachia* to host genome copies was quantified by qPCR using previously developed primers for the *Wolbachia* surface protein (*wsp*) gene and the *Ae. aegypti* Ribosomal protein subunit 17 (*RPS17*) gene ^21^ (Supplementary Table S1).

Total RNA was extracted from three pools of four adult females at 2, 6, and 12 days post-emergence using the Qiazol reagent according to the manufacturer’s instructions (Qiagen). Genomic DNA was removed using the Turbo DNA-free kit (Invitrogen). RNA quality and quantity were evaluated using a BioTek Epoch Microspot plate spectrophotometer. Small RNAs were reverse transcribed with miSCRIPT II RT kit (QIAGEN) using the HiSpec buffer and 0.5 µg of total RNA per reaction. Thermocycling conditions were as per the manufacturer’s instructions. The miSCRIPT RT kit works by poly-adenylating small RNA species present in total RNA. A poly-T primer with a 5’ universal tag is then used for reverse transcription of the small RNAs. Quantitative PCR was performed with a miScript SYBR Green PCR kit (QIAGEN) in a QIAGEN Rotor-Gene Q 2plex using 10 ng of cDNA per reaction. For each small miRNA or 5s rRNA, the small RNA sequence was used for the forward primer sequence (Supplementary Table S1). The miScript Universal Primer (Qiagen) was used as the reverse primer.

### miRNA inhibitor trials

Aag2.*w*AlbB cells were seeded into 12-well plates at a density of 5x10^5^ cells per well, and allowed to adhere. Medium was removed and replaced with a 300µl of serum-free medium containing 100nmol of miRNA inhibitor (GenePharma) in Cellfectin II reagent (Invitrogen). miRNA inhibitors are RNA oligos, chemically modified to bind with high affinity to target miRNAs. Target specificity derives from sequence complementarity, which is 100%. Following incubation at 28°C for three hours, 300µl of medium containing 2% FBS was added to cells. Cells were then grown under normal culturing conditions for three days. DNA and RNA were extracted and used for measuring *w*AlbB density and miRNA inhibition by qPCR and miScript RT-qPCR, respectively, as described above. The effect of inhibition of aae-miR-2b-3p, aae-miR-190-5p, and aae-miR-276b-5p on *w*AlbB density was examined *in vivo* by intrathoracic microinjection of adult females with 125nl of 100µM inhibitor (GenePharma) or a scrambled negative control at 2 days post emergence. As an additional control, mosquitoes were injected with 125nl of phosphate-buffered saline (PBS). Three days after injection, DNA and RNA were extracted and used for measuring *w*AlbB density and miRNA inhibition respectively.

The effect of inhibition of aae-miR-317-3p and aae-miR-2946-3p on mosquito longevity was performed by intrathoracic microinjection of adult females with 125nl of 100µM inhibitor, (GenePharma) or a scrambled negative control (Supplementary Table S1), or PBS at 2 days post emergence. Mosquitoes that survived the 24-hour period following injection were included in the experiment. Three days after injection, inhibition was measured by miScript qPCR as described above. The experiment was performed twice, with 10 to 12 mosquitoes per treatment, per experiment. Data were tested for proportionality using the cox.zph function in R, and analysed using a Cox proportional hazards model, for the effect of treatment while controlling for batch effect, using the survminer package in R. Tests returning a likelihood ratio test *p* value < 0.05 were considered significant. A Kaplan-Meier estimator was used to calculate median survival estimates for each group.

### Quantification and statistical analysis

All statistics were performed in R (version v3.6.2) or GraphPad Prism v. 9.1.2. Significance for differences between treatment and control groups were determined by statistical analyses mentioned in each relevant figure legend. Where data were approximately normally distributed, a parametric test was used. Otherwise a non-parametric test was used.

### Data and code availability

The accession number for the small RNA-Seq dataset reported in this paper is SRA: PRJNA671731. Reviewers can access the data through this link: https://dataview.ncbi.nlm.nih.gov/object/PRJNA671731?reviewer=p9kc0vvesp2t19cduivn46504t.

## Results

### Mosquito rearing and tetracycline clearance

To produce a *Wolbachia*-free line as control, *w*AlbB-transinfected *Ae. aegypti w*AlbB2-F4 (WB2) mosquitoes (males and females) were treated with tetracycline (WB2.tet). We confirmed the presence or absence of *Wolbachia w*AlbB in mosquitoes using qPCR of genomic DNA from a sample of female mosquitoes in three age groups: 2, 6 and 12 days post-emergence. No *Wolbachia* surface protein gene (WSP) product was detected in female WB2.tet mosquitoes. In female WB2 mosquitoes, *w*AlbB density was not significantly different between age groups (ANOVA, *p* > 0.05) (Fig. 1A).

**Figure 1:**
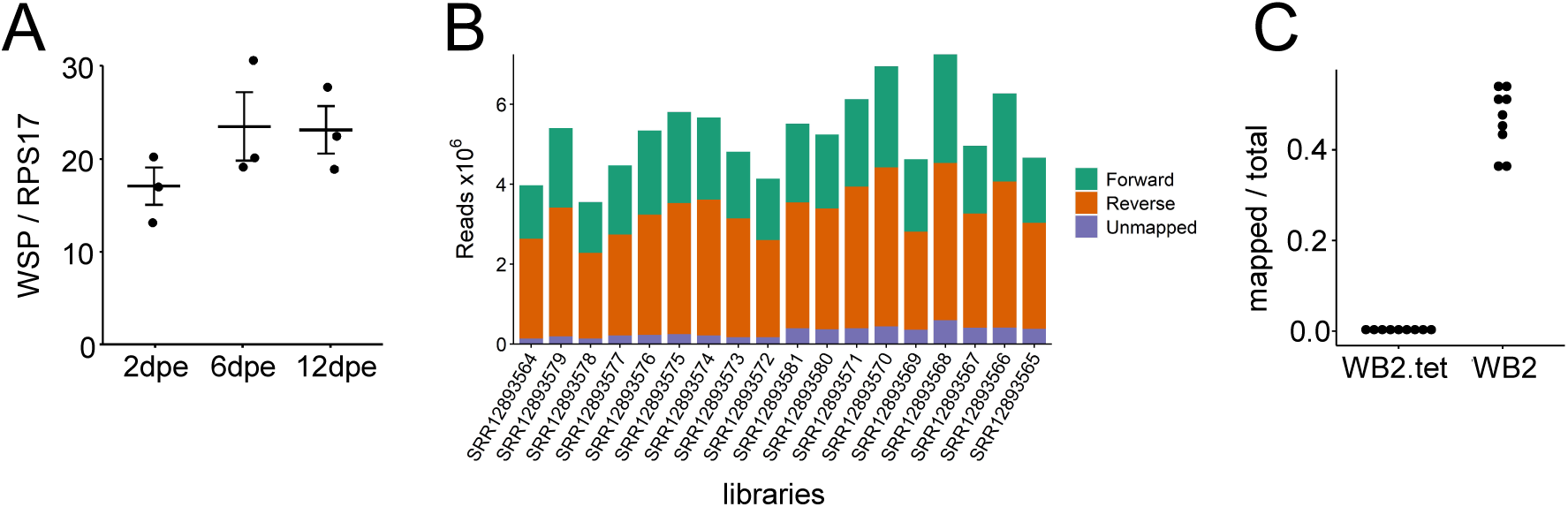
*w*AlbB density in WB2 mosquitoes and results of read mapping. **A)** Relative *w*AlbB density at 2, 6, and 12 dpe. Crossbars represent the mean of three replicates (n=3). Density of *w*AlbB was not significantly different between age groups (ANOVA, *p* > 0.05). Error bars represent SEM. **B)** Number of forward- and reverse-mapped reads to the *Ae. aegypti* AaegL5.0 genome after size selection and quality filtering. **C)** Reads that did not align to the *Ae. aegypti* genome were aligned to the *Wolbachia w*AlbB genome. Each point represents the proportion of non-*Ae. aegypti* reads that mapped to *Wolbachia w*AlbB.

### Small RNA sequencing

To compare the miRNA profiles of WB2 and WB2.tet mosquitoes, we collected mosquitoes at 2, 6 and 12 dpe, from which RNA was extracted for small RNA sequencing (sRNA-Seq). Small RNA sequencing of 18 RNA libraries yielded a total of 94.8 million single-end reads with a Phred score of ≥Q30 after quality trimming and size selection (Supplementary Table S2). After removing sequencing adapters, the read length profile of each library was characteristic of a small RNA fraction, with a sharp peak at 21-23nt corresponding to miRNA and short interfering-RNA classes, and a broader peak at 26-31nt corresponding to PIWI-RNAs (Supplementary Fig. S1). Reads of 18-24nt in length were aligned to the *Ae. aegypti* AaegL5.0 genome, with an alignment rate of between 91.8% and 96.5% per sample (Supplementary Table S2). The total number of aligned 18-24nt reads ranged from 3.4 to 6.6×10^6^ reads per sample (Fig. 1B). Following tetracycline treatment, each mosquito line was reared under identical conditions for five generations. Nonetheless, to ensure that the bacterial composition of each line was consistent, we conducted a metagenomics analysis using reads that did not align to the AaegL5.0 genome. This confirmed that the bacterial composition was congruent between lines (Supplementary Fig. S2). To confirm the absence of *w*AlbB from the WB2.tet samples, reads that did not map to the AaegL5.0 genome were mapped to the *w*AlbB genome. For the *w*AlbB libraries, 3.42% of the reads mapped to the *w*AlbB genome, whereas for the WB2.tet samples, 0.01% mapped, mainly to the 16s and 23s rRNA loci (Fig. 1C). To determine the validity of these mappings, we collected the *w*AlbB-mapped WB2.tet reads and mapped them against the genomes of five bacterial strains that were identified to the genus level in the metagenomic analysis. Of those reads, 68.19% mapped to at least one of the five strains, mainly to the rRNA loci. We therefore concluded that the small fraction of WB2.tet reads mapping to the *w*AlbB genome was likely the result of sequence similarity between bacterial genomes, the intrinsic error-rate associated with mapping short reads with a high sensitivity mapping algorithm, and/or the result of index-hopping during library preparation.

### Annotation and differential expression analysis

Previous annotations of precursor and mature miRNAs in *Ae. aegypti* relied on the scaffold-level AaegL3.0 reference genome that is now superseded ^73^. We annotated miRNAs using the current AaegL5.0 genome, which contains all three complete chromosome assemblies and 2309 scaffolds ^58^. As a result of the improved completeness of the reference genome, 13 miRNAs that were previously reported to be duplicated ^42^ were shown to occur as single copies in the full-length chromosome assemblies and to be absent from the scaffolds. A BLAST analysis of our miRNA sequences indicated that nine other miRNAs had potential duplicates on a scaffold. The AaegL5.0 genome was produced from a pool of 80 individuals, and many of the scaffolds are haplotigs, i.e. the result of alternative haplotypes present in the pool ^58^. To confirm the validity of the duplicates, we aligned the chromosome copy and scaffold copy of each potential duplicate, including 500b up- and downstream flanking regions. All nine ‘chromosome’ copies had >95% similarity with their respective ‘scaffold’ copy across the ∼1100b alignment. We therefore assumed that the scaffolds on which those nine duplicates occurred were haplotigs, and the duplicates were artificial. Since no unique miRNAs were found on a scaffold, we removed the scaffolds from our analyses. Genomic coordinates for 221 mature and 116 precursor miRNA loci within the three full chomosomes of the AaegL5.0 genome are provided in Supplementary Table S3. By comparing our annotations to the AaegL5.0 genome, we determined the genomic context of each miRNA locus with respect to being intergenic, intronic, or exonic, as well as identifying potential miRNA clusters (Fig. 2). miRNAs with expression values greater than five counts-per-million in at least three samples were included in the statistical analysis, resulting in 168 mature miRNAs being examined. A principal components analysis of log_2_ normalised counts showed some separation of WB2 and WB2.tet on the second principal component for the 2 dpe and 6dpe age groups (Fig. 3A). However, separation was greatest between age groups, particularly for WB2 groups, which clearly separated along the first principal component (Fig. 3A). In the 2 dpe group, differential expression was approximately even in terms of up- vs down-regulation, whereas 6 dpe and 12 dpe groups, were skewed toward downregulation (Fig. 3B). For comparison between *Wolbachia*-infected and uninfected mosquitoes, miRNAs with an adjusted p value of < 0.05 and an absolute log_2_ fold-change of > 0.5 were considered differentially expressed. A total of 44 miRNAs met these criteria (Table 1). Log_2_ fold-changes ranged from -2.05 to 1.57 (Table 1). There was limited overlap in miRNAs that showed differential expression between lines (Fig. 3C). Two miRNAs, aae-miR-12-5p and aae-miR-998-5p, were consistently differentially expressed in both 2 dpe and 6 dpe groups (Table 1). A third miRNA, aae-miR-309a-3p, was upregulated at 2 dpe and downregulated at 12 dpe (Table 1). No miRNAs were significantly differentially expressed across all three age groups. Given the apparent lack of consistency between age groups, we compared log_2_ fold-changes of miRNAs that were significantly differentially expressed in one age group against log_2_ fold-changes of the same miRNAs in other age groups - most of which were not significantly differentially expressed in the second age group. All comparisons showed significant correlation (Pearson, *p* < 0.05), suggesting that despite a lack of overlap between age groups in terms of which miRNAs passed the cut-off for differential expression, the broader transcriptional profiles did show some consistency between age groups (Fig. 3D).

**Figure 2:**
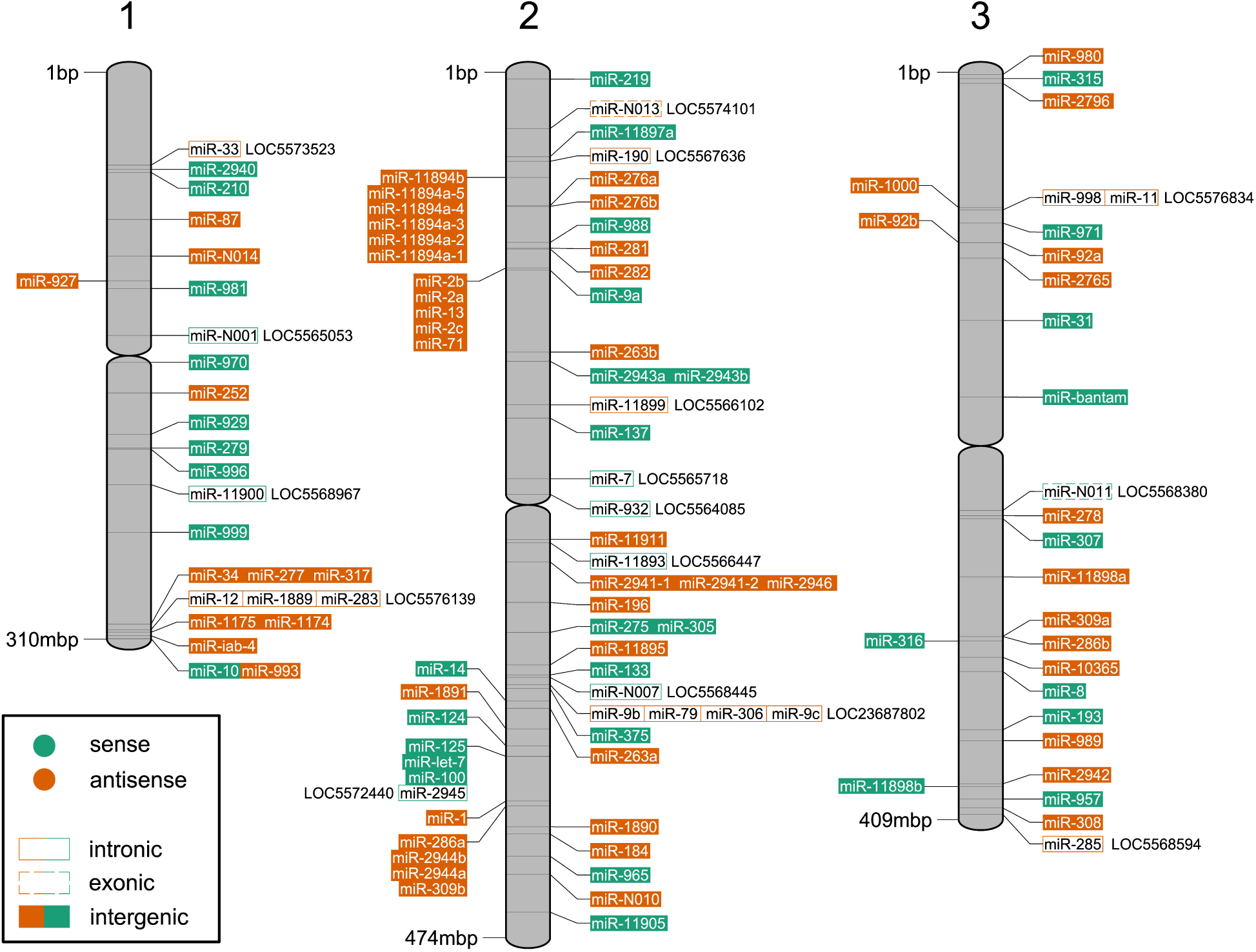
Position of miRNA loci on *Ae. aegypti* chromosomes. Map of the three *Ae. aegypti* chromosomes showing position, strand, and genomic context (i.e. intronic, exonic, or intergenic) of precursor miRNA loci. Gene identifiers with ‘LOC’ prefix indicate parent genes of intronic or exonic miRNA loci.

**Figure 3:**
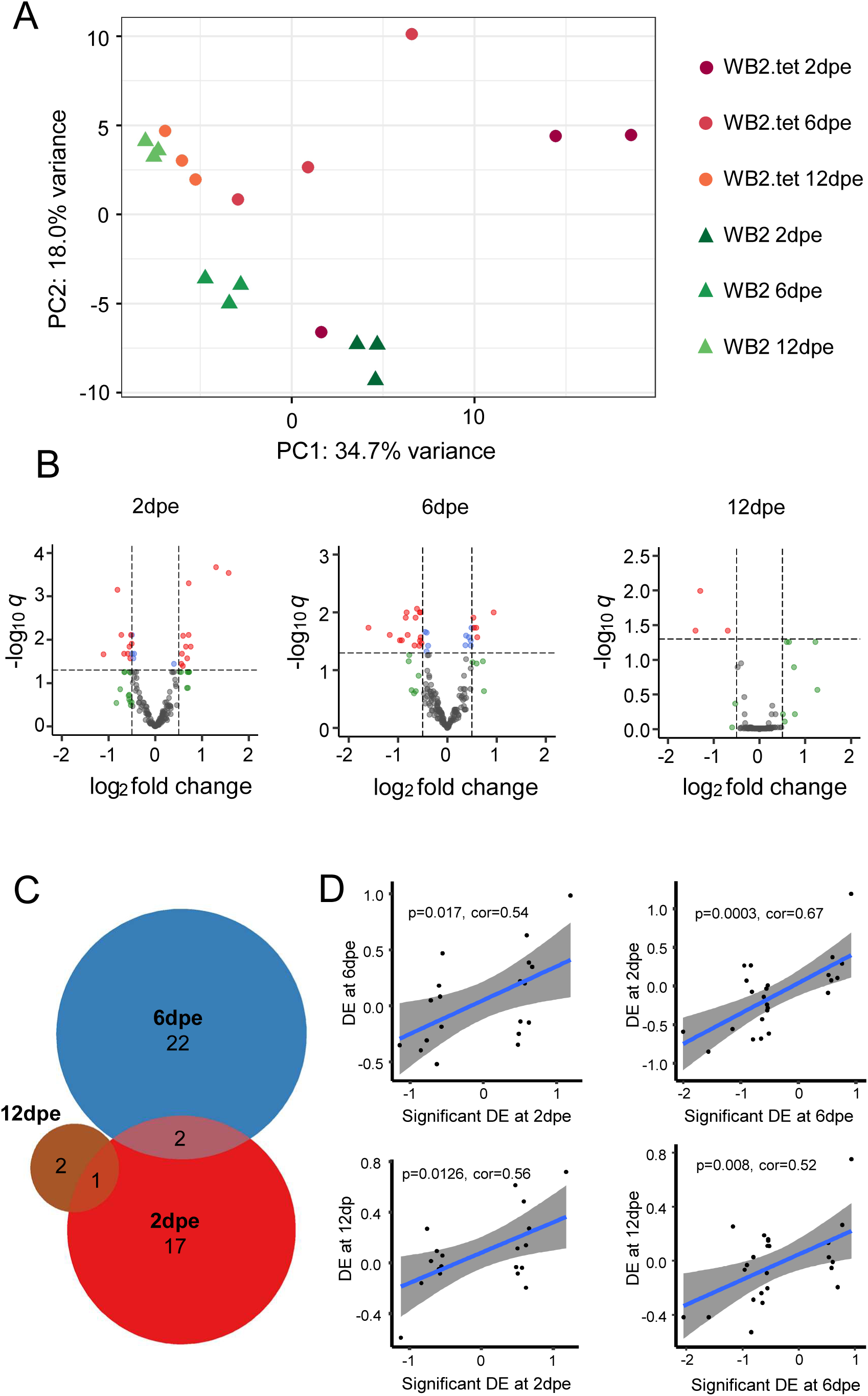
Differentially expressed miRNAs in WB2 vs. WB2.tet mosquitoes. A) Principal components analysis of log_2_ normalised counts of 168 miRNAs. **B)** Volcano plots show differential expression of miRNAs in each age group. Red dots represent miRNAs that meet the cut-off for FDR (*q*) and log_2_ fold-change; green, those that meet log_2_ fold-change cut-off, but not *q* significant; blue, those that meet the *q* significance but not the log_2_ fold-change cut-off; grey, those that meet neither cut-off. **C)** Venn diagram showing overlap in significantly differentially expressed miRNAs between three age groups: 2 dpe, 6 dpe and 12 dpe. Only two miRNAs, miR-12-5p and miR-998-5p were in common between 2 dpe and 6 dpe. A third miRNA, miR-309a-3p, was down-regulated at 2 dpe and up-regulated at 12 dpe. **D)** Pairwise comparisons of log_2_ fold-changes of miRNAs that were significant in one age group (x-axis) against the same miRNAs in another age group (y-axis). All Pearson correlations were significant (p < 0.05).

**Table 1:** Differential expression of 46 miRNAs in WB2 vs WB2.tet mosquitoes in three age groups. Log_2_ fold-changes > 0 represent upregulation in WB2 compared to WB2.tet. log_2_ CPM, counts per million reads mapped to each miRNA given (log_2_). FDR, false discovery rate (Benjamini-Hochberg corrected p-value).

| miRNA | log<sub>2</sub> CPM | log<sub>2</sub> FC | FDR | Age group |
| --- | --- | --- | --- | --- |
| 309a-3p | 4.14 | 1.57 | 2.88E-04 | 2dpe |
| 998-5p | 5.05 | 1.30 | 2.12E-04 | 2dpe |
| 10365-3p | 4.77 | 0.76 | 1.44E-02 | 2dpe |
| 2941-2-3p | 11.74 | 0.72 | 4.98E-04 | 2dpe |
| 11-5p | 8.94 | 0.71 | 7.75E-03 | 2dpe |
| 2765-5p | 5.25 | 0.68 | 2.68E-02 | 2dpe |
| 278-3p | 12.10 | 0.65 | 1.44E-02 | 2dpe |
| 2941-1-3p | 8.94 | 0.59 | 8.11E-03 | 2dpe |
| 965-5p | 5.42 | 0.59 | 4.03E-02 | 2dpe |
| 989-5p | 5.39 | 0.57 | 2.10E-02 | 2dpe |
| 1174-5p | 5.86 | 0.56 | 3.60E-02 | 2dpe |
| 100-3p | 8.14 | -0.50 | 1.23E-02 | 2dpe |
| 1000-5p | 7.54 | -0.52 | 7.75E-03 | 2dpe |
| 279-3p | 12.73 | -0.54 | 2.68E-02 | 2dpe |
| 10365-5p | 8.89 | -0.55 | 1.44E-02 | 2dpe |
| 12-5p | 9.64 | -0.58 | 2.10E-02 | 2dpe |
| let-7-5p | 13.45 | -0.67 | 2.10E-02 | 2dpe |
| 31-3p | 6.41 | -0.73 | 7.75E-03 | 2dpe |
| 276b-5p | 6.37 | -0.81 | 7.06E-04 | 2dpe |
| 193-3p | 2.40 | -1.11 | 2.17E-02 | 2dpe |
| 998-5p | 5.05 | 0.94 | 9.98E-03 | 6dpe |
| 308-5p | 9.39 | 0.78 | 5.72E-06 | 6dpe |
| 184-3p | 14.72 | 0.70 | 9.95E-05 | 6dpe |
| 13-5p | 6.83 | 0.61 | 2.69E-02 | 6dpe |
| 932-3p | 6.60 | 0.59 | 1.84E-02 | 6dpe |
| 100-5p | 15.53 | 0.54 | 1.84E-02 | 6dpe |
| 10-5p | 13.41 | 0.53 | 1.23E-02 | 6dpe |
| 12-5p | 9.64 | -0.52 | 3.39E-02 | 6dpe |
| 13-3p | 11.15 | -0.54 | 2.69E-02 | 6dpe |
| 277-3p | 14.77 | -0.54 | 9.98E-03 | 6dpe |
| 9a-3p | 5.22 | -0.55 | 3.07E-02 | 6dpe |
| 92a-3p | 8.18 | -0.56 | 9.98E-03 | 6dpe |
| 965-3p | 5.26 | -0.57 | 3.77E-02 | 6dpe |
| 1175-3p | 10.50 | -0.62 | 8.62E-03 | 6dpe |
| 2940-5p | 13.38 | -0.64 | 1.23E-02 | 6dpe |
| 996-3p | 6.86 | -0.66 | 3.73E-02 | 6dpe |
| 190-5p | 11.75 | -0.80 | 2.45E-02 | 6dpe |
| 2946-3p | 9.98 | -0.83 | 9.98E-03 | 6dpe |
| 286b-3p | 5.33 | -0.84 | 1.23E-02 | 6dpe |
| 219-5p | 3.13 | -0.92 | 3.03E-02 | 6dpe |
| 980-5p | 4.52 | -0.96 | 3.03E-02 | 6dpe |
| 2944a-5p | 2.84 | -1.17 | 2.45E-02 | 6dpe |
| 2944b-3p | 2.38 | -1.60 | 1.84E-02 | 6dpe |
| 2942-3p | 3.84 | -2.05 | 1.48E-06 | 6dpe |
| 309a-3p | 4.14 | -0.69 | 3.79E-02 | 12dpe |
| 2941-1-5p | 3.39 | -1.29 | 1.01E-02 | 12dpe |
| 196-3p | 2.93 | -1.40 | 3.79E-02 | 12dpe |

To validate differential expression of miRNAs between lines, we used a new generation of mosquitoes and RT-qPCR to test ten miRNAs that showed relatively high log_2_ fold-changes in the original small RNA-Seq analysis (Supplementary Table S4). One miRNA was too lowly expressed in whole mosquitoes to amplify sufficiently via RT-qPCR, and was therefore excluded from analysis. Seven miRNAs (190-5p, 276b-5p, 2940-5p, 2941-1-3p, 2946-3p, 309a-3p, and 31-3p 2dpe) showed a consistent trend with small RNA-Seq in at least two out of three age groups, while a further two miRNAs (308-5p, and 184-3p) showed a direction of change that was inconsistent with small RNA-Seq (Fig. 4).

**Figure 4:**
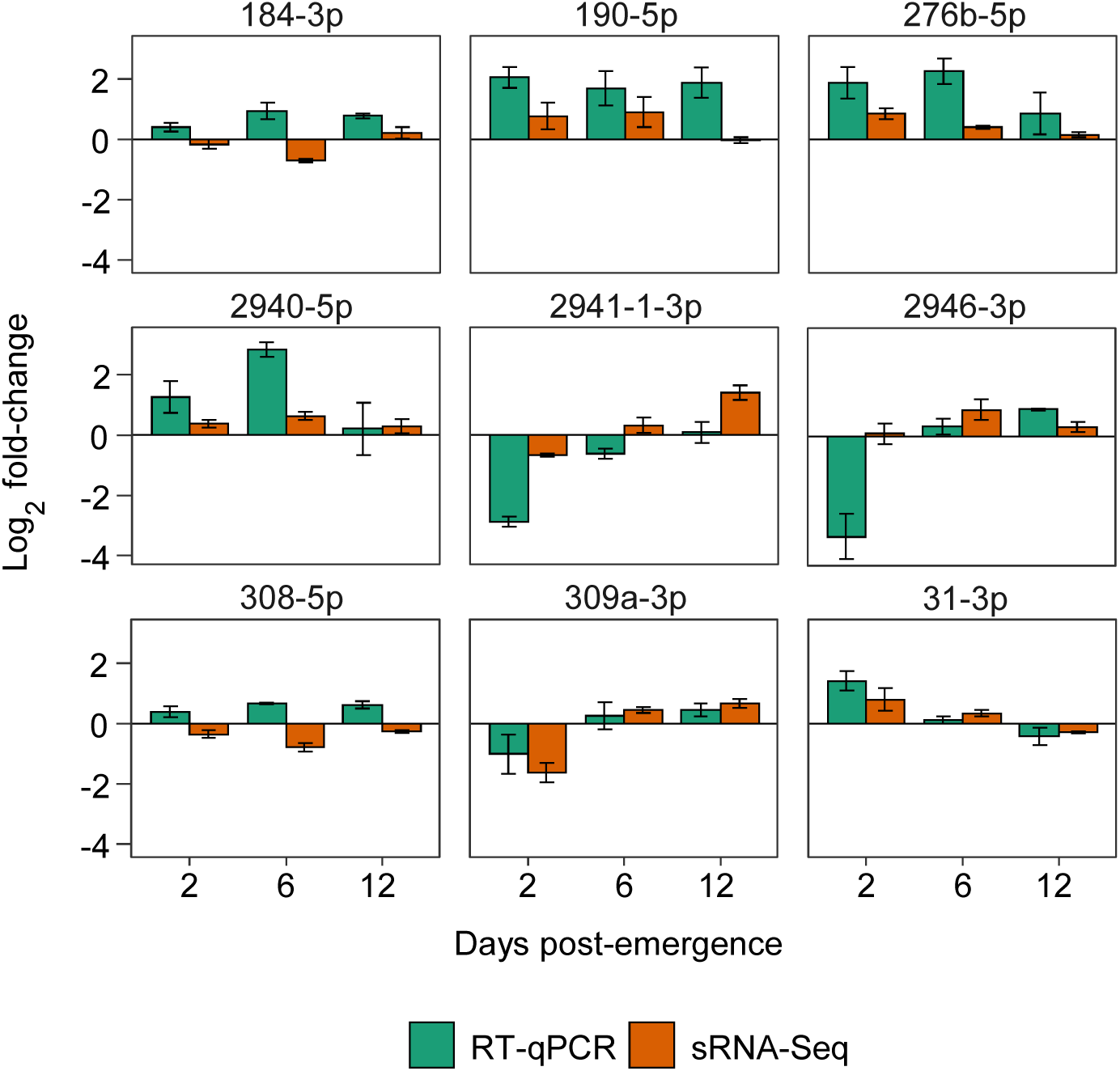
Validation of selected differentially expressed miRNAs. Plots show log_2_ fold changes of a selection of miRNAs according to small RNA-Seq and RT-qPCR at 2, 6, and 12 days post-emergence using 5S RNA for normalization of data. Bars are averages of three biological replicates. Error bars show standard error of the mean.

We next investigated the effect of age on miRNA expression. Generally, there was greater differential expression between age groups than between lines (Fig. 3A), therefore, for classifying miRNAs as being differentially expressed, we increased the log_2_ fold-change threshold from > 0.5 to > 1. A total of 34 miRNAs met these criteria (Table 2). Of these, 14 showed age-related differential expression in both lines (2940-5p, 34-5p, 317-3p, 11894b-3p, 11894a-4-3p, 11899-5p, 2946-3p, 1-5p, 1890-3p, 2765-5p, 309a-3p, 193-5p, 2765-5p, and 193-5p) (Fig. 5A, Table 2). Log_2_ fold-changes ranged from -2.61 to 1.56 in WB2.tet groups, and -2.70 to 3.47 in WB2 groups (Table 2). A total of 15 miRNAs that were differentially expressed with age were also differentially expressed between lines (277-3p, 2941-1-3p, 2941-2-3p, 2946-3p, 2765-5p, 309a-3p, 286b-3p, 193-3p, 2942-3p, 279-3p, 996-3p, 276b-5p, 2941-1-5p, 196-3p, and 989-5p) (Table 1 and 2).

**Figure 5:**
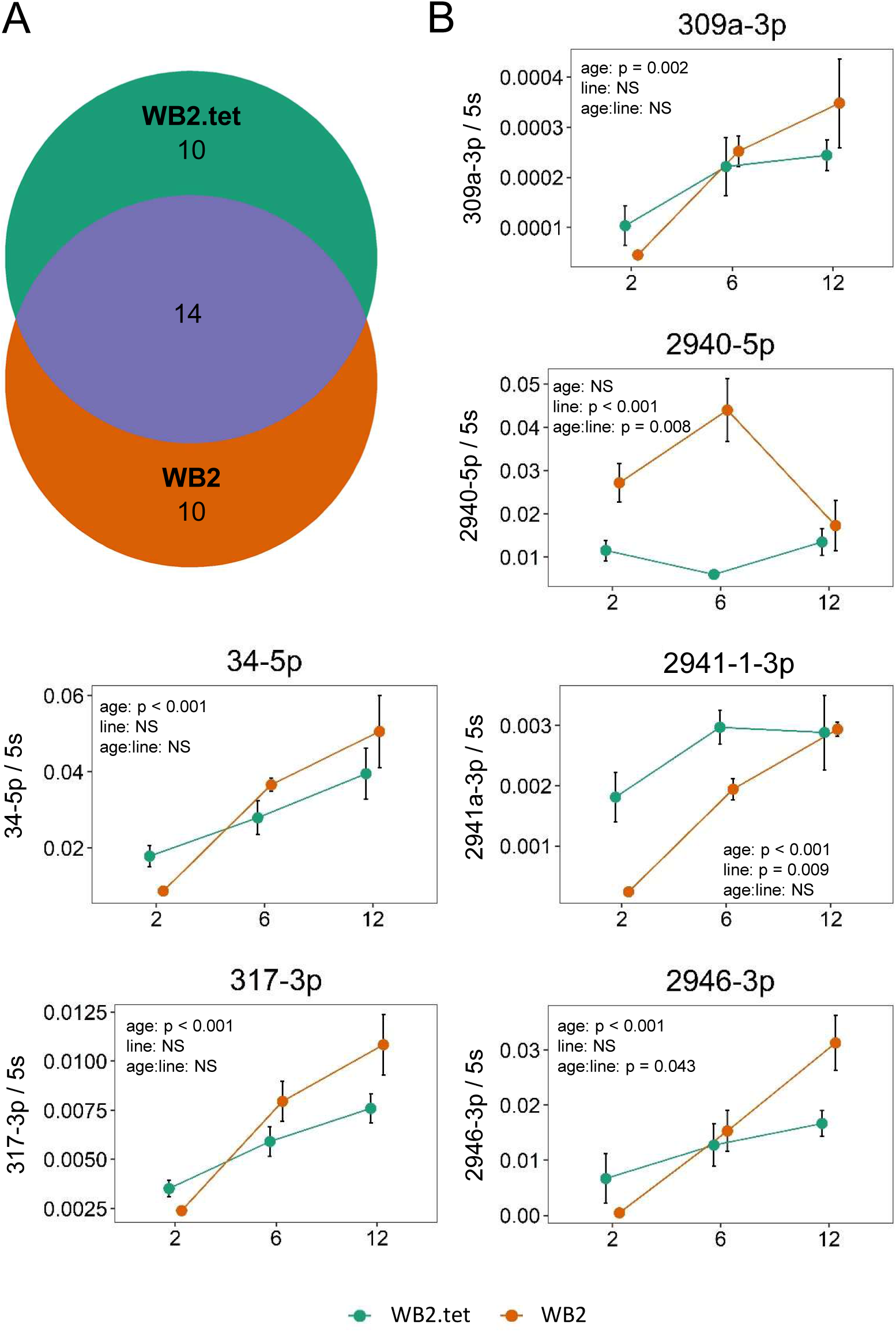
Expression of *Ae. aegypti* miRNAs increasing with age. **A)** Venn diagram showing overlap in significantly differentially expressed miRNAs between two mosquito lines. 14 out of 34 miRNAs (41%) were differentially expressed with age in both WB2 and WB2.tet mosquitoes. **B)** Graphs showing results of RT-qPCR of six miRNAs that showed significant differential expression between age groups according to RNA-Seq. Points are averages of three biological replicates. Error bars represent standard error of the mean. Significant difference between means was estimated using a factorial ANOVA (p < 0.05). NS, not significant.

**Table 2:**
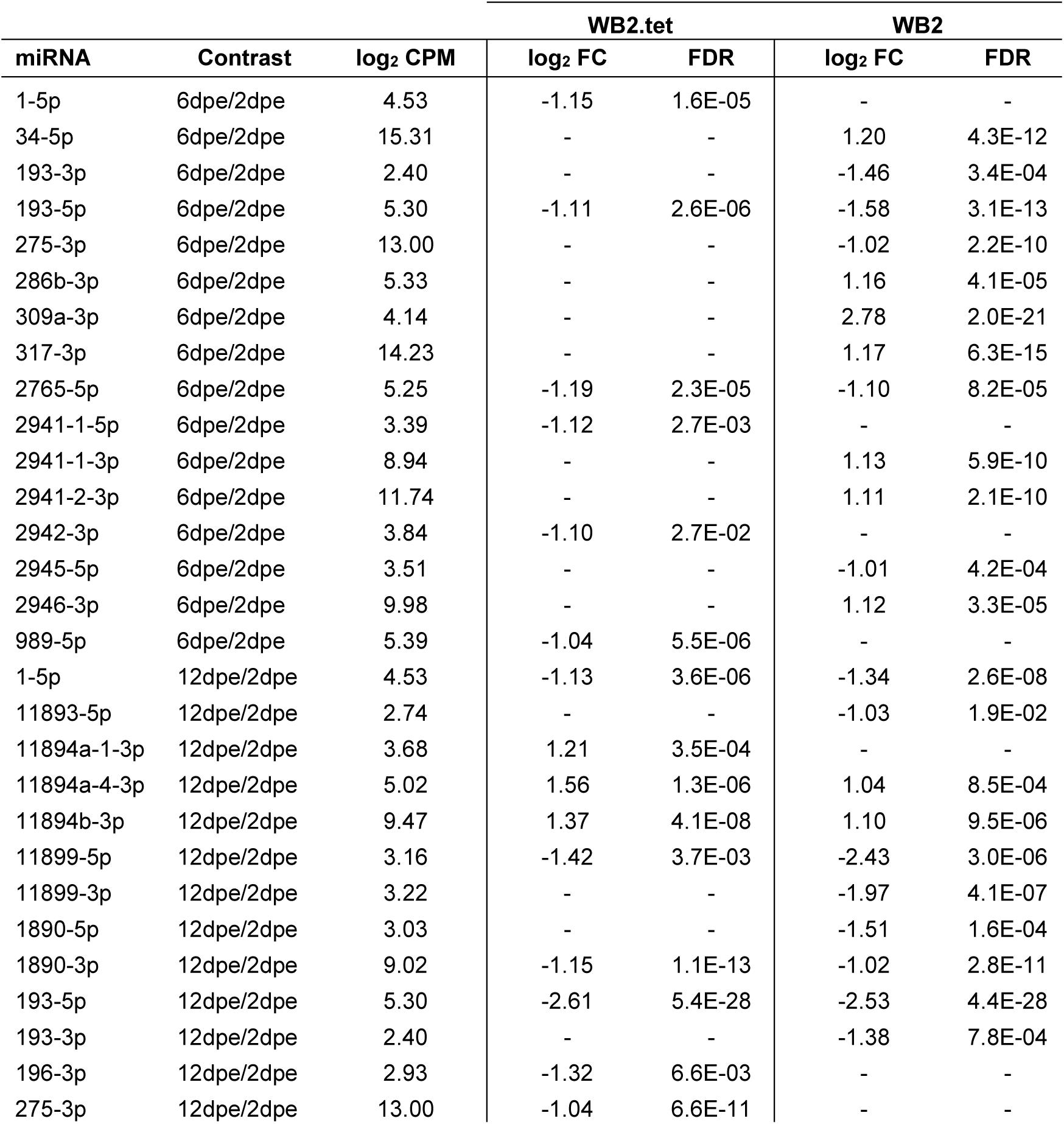
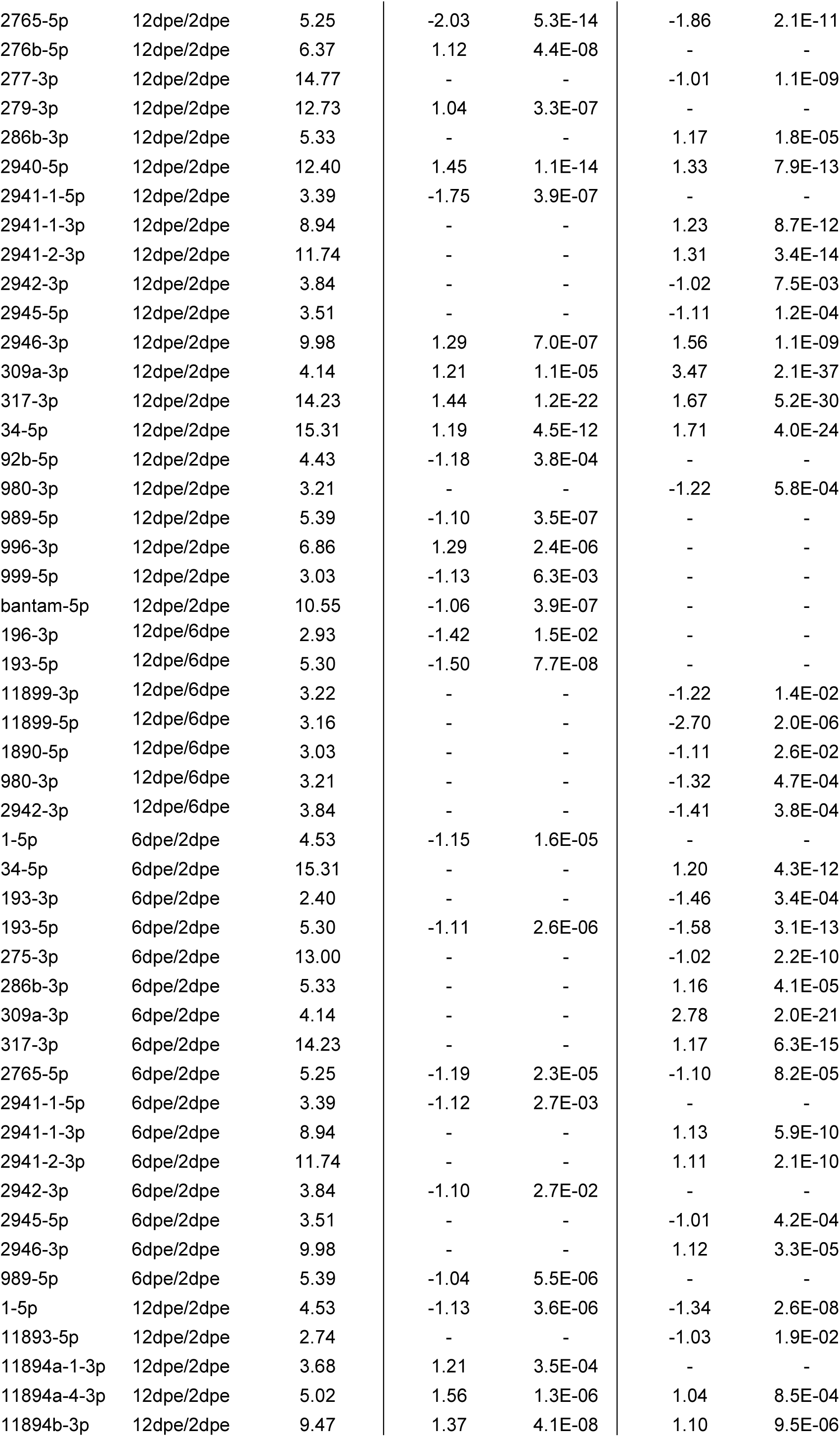
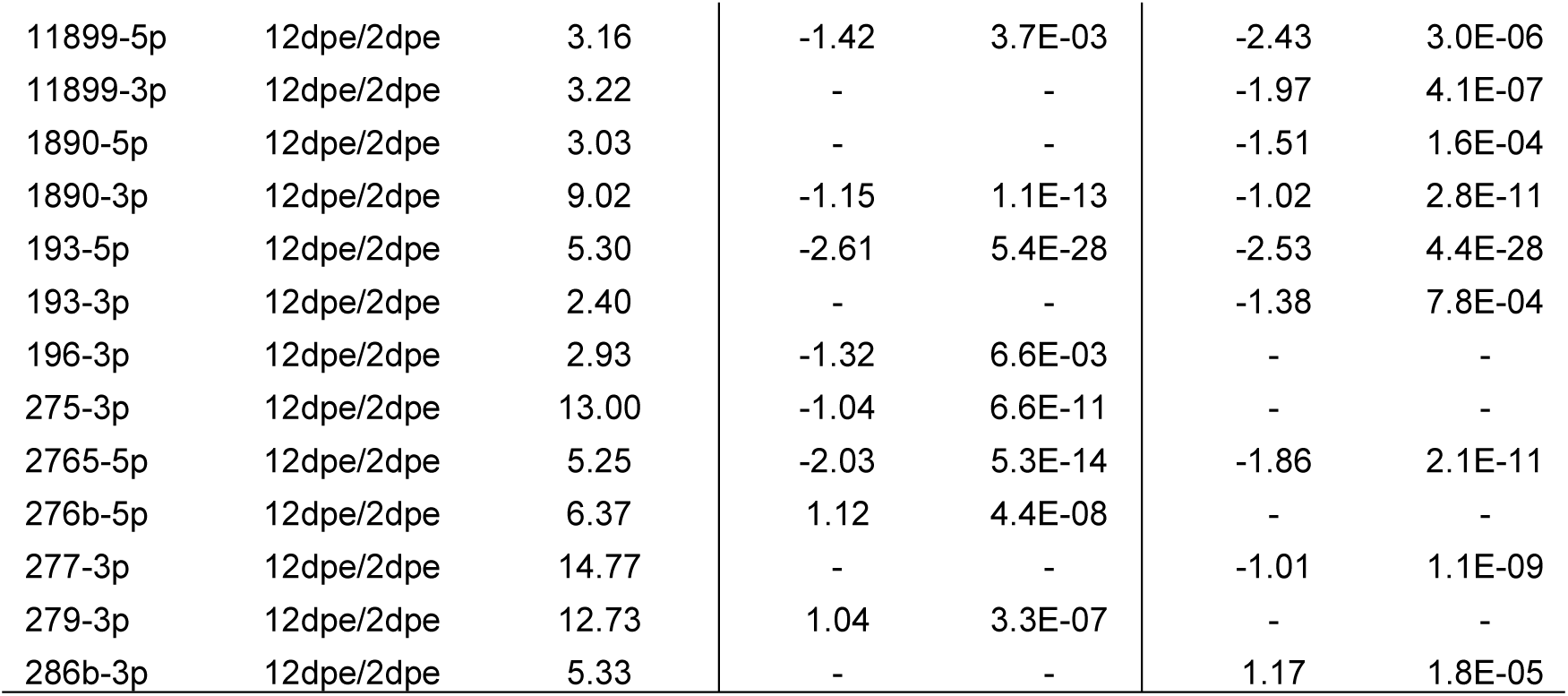
Differential expression of 31 miRNAs in older versus younger age groups in each mosquito line. Fold-changes > 0 represent upregulation in the older age group. log_2_ CPM, counts per million reads mapped to each miRNA (log_2_).

To validate differential expression of miRNAs between age groups, we used a new generation of mosquitoes and RT-qPCR to test ten miRNAs that showed relatively high log_2_ fold-changes and/or showed differential expression in both lines in the original sRNA-Seq (Supplementary Table S4). Four miRNAs were too lowly expressed in whole mosquitoes to amplify sufficiently via RT-qPCR, and were therefore excluded from analysis. Of the six remaining miRNAs, five were significantly upregulated with age (ANOVA, *p* < 0.05) (Fig. 5B). miR-2941-1-3p and 2940-5p were both significantly different between lines, while 2946-3p and 2940-5p both indicated a significant interaction between age and line (ANOVA, *p* < 0.05) (Fig. 5B). All miRNAs were initially lower in WB2 than in WB2.tet, but increased at a greater rate to become even with or higher than WB2.tet (Fig. 5B). miR-2940 showed differential expression with age in WB2 mosquitoes, but not in WB2.tet (Fig. 5B).

### Target prediction of differentially expressed miRNAs

To determine if the expression of the predicted targets of the differentially expressed miRNAs could be affected by age, we selected three highly confident targets of aae-miR-317-3p, and aae-miR-2946-3p (Table 3). RNA extracted from WB2 and WB2.tet mosquitoes collected at 2, 6, and 12 dpe was subjected to RT-qPCR. aae-miR-317-3p and aae-miR-2946-3p were upregulated in mosquitoes with age. RT-qPCR results revealed differential expression of most of the predicted target genes of the two miRNAs with age (ANOVA, *p* < 0.05) (Fig. 6). For the targets of 317-3p, AAEL010793 and AAEL010508 were significantly downregulated at 6 and 12 dpe compared to 2 dpe, while the transcript levels of AAEL006171 steadily increased over time (Fig. 6). For 2946-3p targets, AAEL006095 did not change over time, AAEL006113 declined at 6 dpe but increased at 12 dpe, and AAEL003402 significantly increased at 6 dpe but returned to the same levels as those of 2 dpe at 12 dpe (Fig. 6). Overall, these results showed that the predicted targets of the differentially expressed miRNAs also responded to age. However, these results are only correlational and direct regulation/interaction of the miRNA-target combinations need to be experimentally validated.

**Figure 6:**
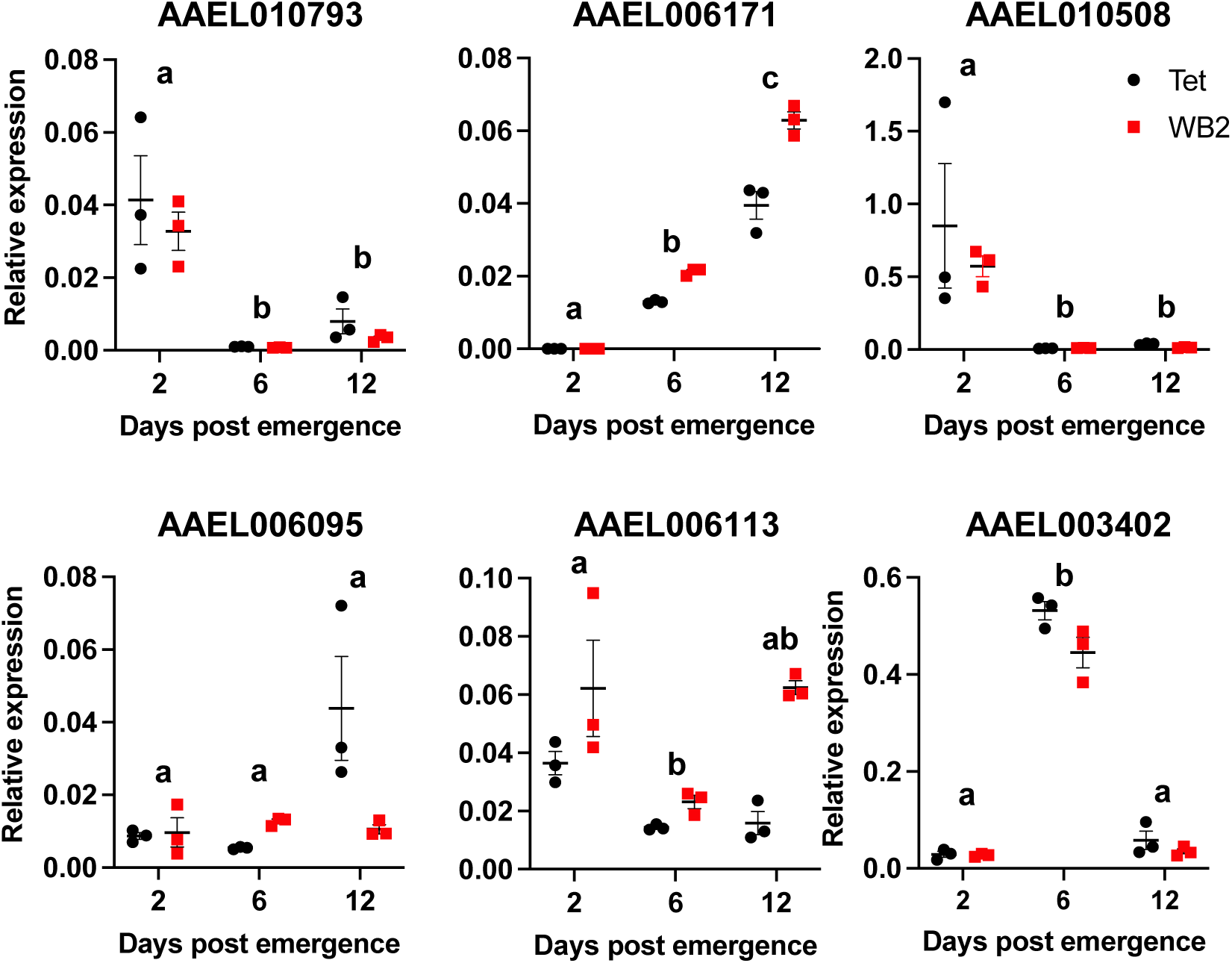
Differential expression of selected potential targets of aae-miR-317-3p (top row) and aae-miR-2946-3p (bottom row) that were differentially expressed with mosquito age. RT-qPCR analysis of RNA extracted from mosquitoes at 2, 6, and 12 dpe using primers to the target genes and *rps17* as the normalizing gene. Two-way ANOVA with post-hoc multiple comparisons was carried out to compare the expression levels between different time points. Different letters on top of each time point represent statistically significant differences.

**Table 3:** High confidence predicted targets of aae-miR-317-3p, and aae-miR-2946-3p. These predicted targets were subjected to RT-qPCR to determine if their expression could be affected by age (Supplementary Fig. S4).

| miRNA | VectorBase accession no. | No. of predicted binding sites | Predicted target gene |
| --- | --- | --- | --- |
| **317-3p** | AAEL010793 | 7 | F-box/leucine rich repeat protein |
| **317-3p** | AAEL006171 | 11 | n-myc downstream regulated |
| **317-3p** | AAEL010508 | 11 | vacuolar protein sorting-associated protein (vps13) |
| **2946-3p** | AAEL006095 | 5 | Gelsolin precursor |
| **2946-3p** | AAEL006113 | 7 | cystinosin |
| **2946-3p** | AAEL003402 | 9 | sphingomyelin phosphodiesterase |

### Functional analysis of differentially expressed miRNAs

We tested the effect of miRNA inhibition on *w*AlbB density in a transinfected Aag2.*w*AlbB cell line by inhibiting two miRNAs, aae-miR-190-5p, and aae-miR-276b-5p, that were among the most consistently upregulated in the WB2 versus WB2.tet mosquitoes according to small RNA-Seq and RT-qPCR. Inhibition was done by transfecting the specific inhibitors (reverse complementary short RNAs) of the miRNAs into Aag2.*w*AlbB cells, and was validated by RT-qPCR three days after transfection. No increase in density was observed in the aae-miR-190-5p, or aae-miR-276b-5p inhibited cells (Fig. 7).

**Figure 7:**
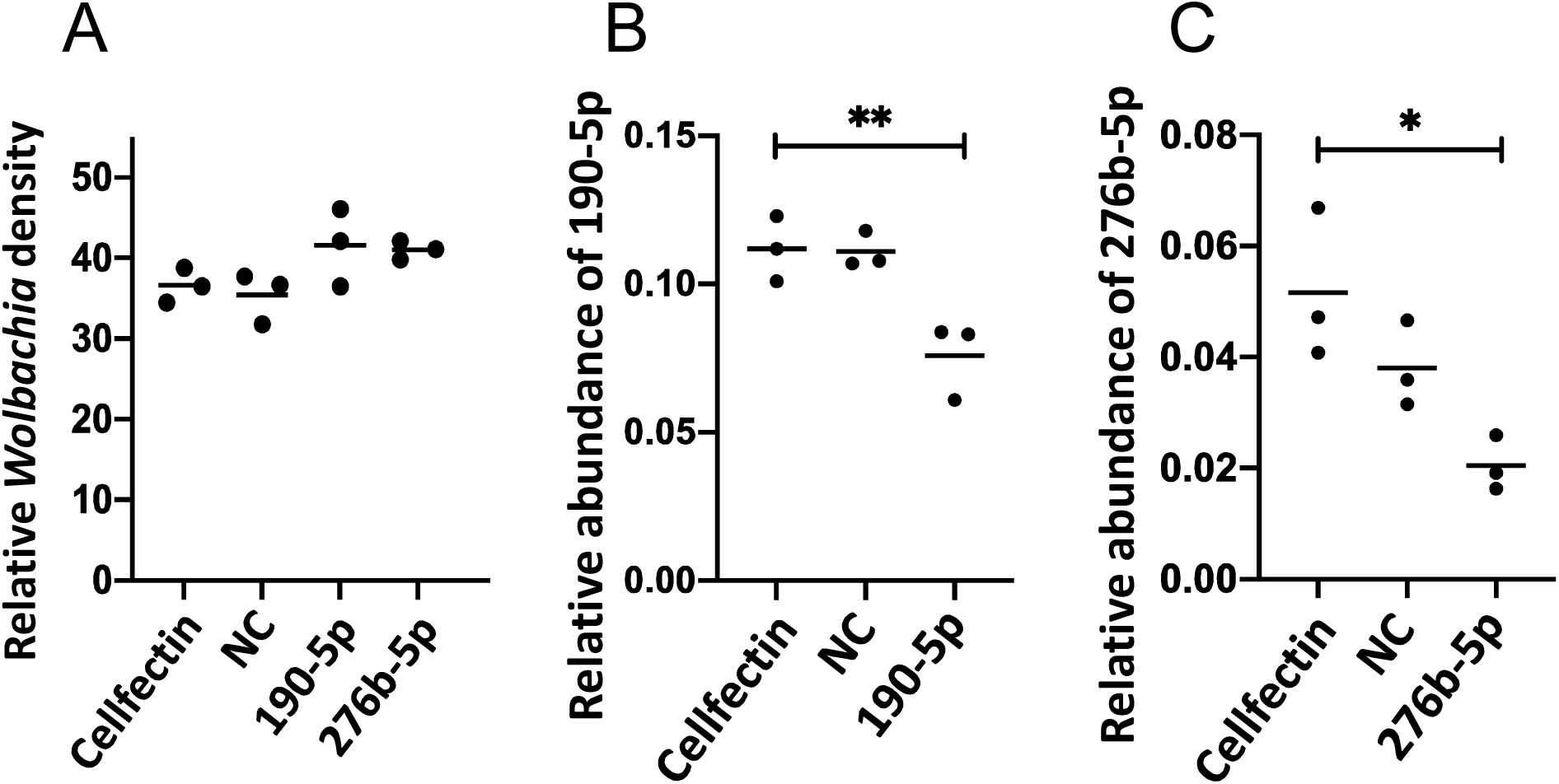
*w*AlbB density in miRNA inhibited Aag2.*w*AlbB cells. **A)** *w*AlbB density of Aag2.*w*AlbB cells following transfection with specific miRNA inhibitors, random negative control (NC) or Cellfectin only. **B and C)** Validation of inhibition of miR-190-5p, and miR-276b-5p by RT-qPCR in the samples used in A. Cross-bars represent the mean of three replicates. Asterisks indicate the level of statistical significance determined using to ANOVA. *, *p* < 0.05; **, *p* < 0.01.

We carried out a small-scale test of the effect of miRNA inhibition on the longevity of WB2 female mosquitoes by inhibiting two miRNAs, miR-317-3p, and miR-2946-3p, that were found to increase with age to a greater degree in WB2 mosquitoes compared to WB2.tet. Three days after miRNA inhibition, miR-317-3p and miR-2946-3p levels were significantly lower (ANOVA, Tukey’s HSD, *p* < 0.001) (Figs. 8A-D). Inhibition did not significantly affect longevity in WB2.tet mosquitoes relative to controls, but in WB2 mosquitoes a significant decrease in longevity was observed in response to inhibition of miR-2946-3p and miR-317-3p compared to negative control (Cox proportional hazard, *p* < 0.0001, and *p* = 0.130, respectively) (Figs. 8E & F). Following injection, the median survival according to a Kaplan-Meier estimator was 26.5 days for miR-2946-3p-inhibited mosquitoes and 35.5 days for the miR-317-3p-inhibited group. A survival estimate was not obtained for the NC and PBS control groups, as not enough deaths occurred within the 40-day period. Longevity between PBS-control mosquitoes did not differ significantly between WB2.tet and WB2 mosquitoes.

**Figure 8:**
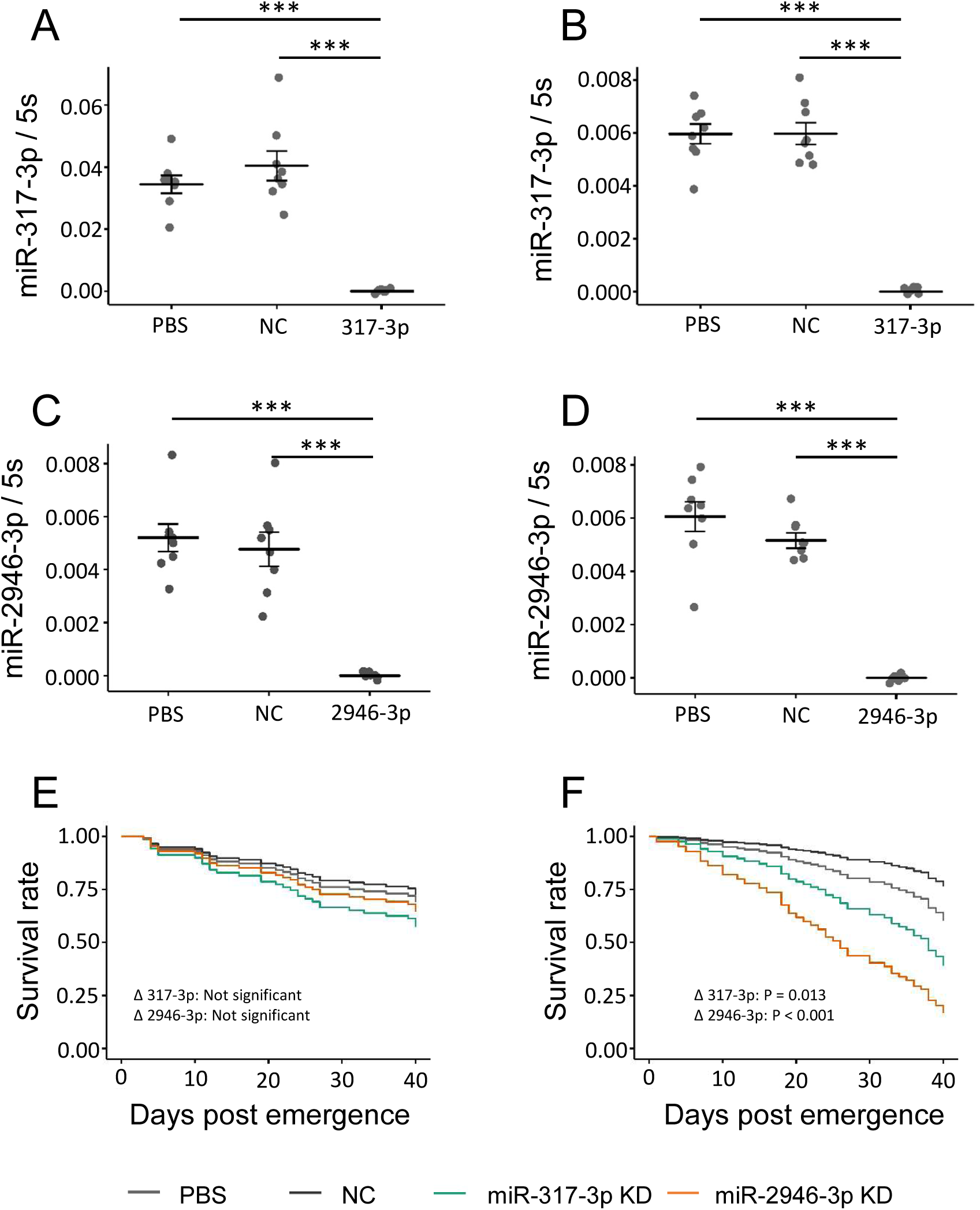
inhibition of miR-317-3p and miR-2946-3p and its effect on lifespan of adult *Ae. aegypti* females. **A and B)** Effect of miRNA inhibition on relative expression of miR-317-3p in WB2.tet and WB2 mosquitoes, respectively. **C and D)** Effect of miRNA inhibition on relative expression of miR-2946-3p in WB2.tet and WB2 mosquitoes respectively. **E and F)** Effect of miRNA inhibition on longevity of WB2.tet and WB2 mosquitoes, respectively. KD, knock down. WB2.tet survival was not significantly affected by miRNA inhibition. WB2 survival was significantly affected by inhibition of miR-2946-3p (Cox ph, *p* < 0.0001) and miR-317-3p (Cox ph, *p* = 0.013).

### Small RNA-Seq reads originating from *w*AlbB

For each age group, WB2 small RNA-Seq read files were concatenated to produce a single read file for each age group. These were filtered on the basis of read length so that only reads 18-24 nt long were retained. Reads were then aligned to the *w*AlbB reference genome, resulting in 769,088, 691,639, and 748,995 aligned reads for 2 dpe, 6 dpe, and 12 dpe groups, respectively. A total of 18 regions of high coverage (greater than 2000×) were identified, with most being present in more than one age group (Supplementary Table S5). Ten peaks corresponded to protein-coding regions, four to intergenic regions, two to tRNAs, one peak corresponded to 5S rRNA, and one to a pseudogene (Supplementary Table S5). Two intergenic regions showed a pattern of read-coverage reminiscent of precursor miRNA loci, and were predicted to form stem-loop structures (Figs. S3-5A). Read-depth of these regions ranged from 3,200 to 16,000× per age group for the concatenated read-files (Figs. S3 and S4). Small-RNA molecules are known to be produced by *w*MelPop-CLA *Wolbachia* ^74^. Sequence alignment of those reported previously ^74^ and the predicted stem-loop structures reported herein showed low sequence similarity (Supplementary Fig. S5). As a control, we concatenated all three replicate fastq files from WB2.tet 2 dpe and mapped reads to the *w*AlbB genome, then counted reads aligned to peaks 1 and 9 that form stem-loop structures found in the WB2 samples. We found only three reads that mapped to peak 1 (compared to ∼10,000 reads mapped in the WB2 samples) and no reads mapped to peak 9.

## Discussion

Using deep sequencing of small RNAs, we identified 116 precursor and 221 mature *Ae. aegypti* miRNA loci, and provide the first chromosomal mapping of these miRNAs. We detected limited differential expression of miRNAs in response to a persistent *w*AlbB-infection in WB2 mosquitoes. We report significant differential expression of 34 mature miRNAs in response to ageing. Further, we identify potential target genes of two miRNAs that were upregulated with age, 317-3p and 2946-3p, and point to a potential link between these miRNAs and longevity. There was limited overlap between age groups in terms of significant differential expression of miRNAs in response to *w*AlbB. However, when miRNAs that were significantly differentially expressed in one age group were compared to the same miRNAs in other age groups, they showed significant positive correlation in log_2_ fold-change, suggesting some degree of consistency in their expression profiles. However, the effect of *w*AlbB on miRNA expression was weak, while the effect of age was robust, and showed reasonable concordance between lines.

We validated the small RNA-Seq results using RT-qPCR by assessing nine *w*AlbB-modulated, and six age-modulated miRNAs that showed relatively high differential expression in small RNA-Seq. For *w*AlbB-modulated miRNAs, the degree of concordance between small RNA-Seq and RT-qPCR was satisfactory, with seven of the nine miRNAs showing agreement between the two methods in at least two age groups. Of the ten age-modulated miRNAs that we tried to validate, four were too lowly expressed in whole mosquitoes to allow sufficient amplification by RT-qPCR. These were all miRNAs that were downregulated with age in the small RNA-Seq analysis (1-5p, 193-5p, 2765-5p, and 2942-3p). Conversely, five of the six validated miRNAs that significantly increased with age were among the most highly expressed in our libraries (34-5p, 317-3p, 2941-1-3p, 2940-5p, and 2946-3p). Another feature of the age-induced miRNAs was that the effect of age was generally more pronounced in the WB2 mosquitoes. Expression of these miRNAs in the WB2 line was significantly lower compared to WB2.tet at 2 dpe for five of the six miRNAs validated by RT-qPCR. Expression in WB2 then increased more dramatically than in WB2.tet, so that by 6 dpe and 12 dpe expression was the same or higher in WB2 mosquitoes. The miR-2940-5p expression pattern was different; there was a significant peak of expression in WB2 mosquitoes at 6 dpe followed by a drop at 12 dpe, whereas in WB2.tet, expression did not change with age.

Some miRNAs form clusters in which individual hairpins are co-transcribed ^75^. The *Ae. aegypti* genome contains a number of conserved miRNA clusters. One example is the miR-2941-1/2941-2/2946 cluster, located in an intron of the putative transcription factor AAEL009263 ^14^. The cluster is synapomorphic to *Aedes* mosquitoes, occurring in both *Ae. aegypti* and *Ae. albopictus*, but absent from other *Culicids* and *Anophelines* ^14, 76^. Our RT-qPCR results for miR-2941-1-3p and miR-2946-3p suggested co-transcription of the two miRNA, with their dramatic suppression in *w*AlbB-infected mosquitoes at 2 dpe, and consistent increase with age. The seed regions of miR-2941-1-3p and miR-2946-3p are identical, while outside of the seed region, nucleotide identity is 25%. Both miR-2941-1-3p and miR-2946-3p are involved in embryonic development in *Ae. aegypti* and *Ae. albopictus* ^76, 77^, and have been observed to increase in response to *w*MelPop infection in *Ae. aegypti* cells ^78^.

miR-34-5p and miR-317-3p belong to the miR-317/277/34 cluster and both increased in abundance with age, which is consistent with reports in *Drosophila* ^79^. miR-277 did not increase with age in either line, which is also consistent with previous observations, and possibly due to a reduced efficiency in the processing of miR-277 by the microprocessor complex ^79^. In *Drosophila*, expression of the 317/277/34 cluster is known to be controlled by the steroid hormone ecdysone ^79^. miR-34-5p has been shown to affect lifespan in flies by modulating branched-chain amino acid catabolism ^80^, and the ecdysone signalling pathway via downregulation of the *E74A* isoform of the transcription factor *Eip74EF* ^79–81^. Furthermore, loss of function mutation in miR-34 causes dramatic life-shortening and neurodegeneration in *Drosophila* ^81^. miR-34 and miR-277 are also involved in innate immunity by regulating Toll signalling in *Drosophila*, with miR-34 promoting Toll signalling and miR-277 having an inhibitory effect ^82^. In *Anopheline* mosquitoes, miR-34 is induced by *Plasmodium* infection, and in *Ae. aegypti* Aag2 cells it is repressed by *w*MelPop infection ^78, 83^.

In addition to the miR-2941-1/2941-2/2946 and miR-317/277/34 clusters, *w*MelPop has been shown to modulate several other miRNAs that were differentially expressed in our WB2 mosquitoes. Our data showed upregulation of miR-190-5p in the WB2 line. In contrast, *w*MelPop was reported to suppress miR-190 in cytoplasmic fractions of *Ae. aegypti* cells ^78^. miR-190 has also been reported to be upregulated in response to CHIKV infection in *Ae. albopictus* ^41^. The ovary-specific miR-309a-3p controls ovarian development and is upregulated by *w*MelPop in *Ae. aegypti* mosquitoes ^43, 84^. We observed an initial suppression of miR-309a-3p in WB2 mosquitoes at 2 dpe, then an upregulation at 6 dpe and 12 dpe, as well as a consistent increase with age in both lines, most strikingly in WB2.

Some differentially expressed miRNAs reported here showed a consistent trend of expression in response to *w*AlbB across all age groups, while for others the effect was dependant on age. Our data also showed that some previously documented age-related miRNAs increased significantly with age in a manner that was affected by *w*AlbB infection. This raises the question of whether *w*AlbB affects the longevity of *Ae. aegypti* via modulation of certain miRNAs. In its native host *Ae. albopictus*, *w*AlbB has been found to decrease longevity in males, but not in females ^85^. In *Ae. aegypti*, the results are less clear. In one study *w*AlbB- infected females exhibited reduced longevity compared to uninfected controls ^15^. However, it is reported elsewhere that *w*AlbB increased longevity following blood feeding ^28^. We investigated the effect of two miRNAs; miR-317-3p and miR-2946-3p, that were significantly upregulated with age and between lines, being higher in WB2 than WB2.tet. Neither miRNA had an effect on longevity in WB2.tet mosquitoes, whereas in WB2 mosquitoes, inhibition was associated with decreased survival. Further, when we compared longevity between WB2.tet and WB2 control groups there was no difference. We hypothesise that the increase in miR-2946-3p and miR-317-3p that we observed with age may form part of a response by *Ae. aegypti* that offsets the fitness cost imposed by *w*AlbB; although to make a more concrete conclusion on the effect of the miRNAs on longevity, this experiment needs to be repeated with larger cohorts of mosquitoes. Additionally, it is possible that the combined stress of *Wolbachia* infection together with the inhibition of any miRNA that is expressed during the adult life span could affect survival. Therefore, to demonstrate a specific role for miR-2946-3p and miR-317-3p in longevity of *w*AlbB-infected mosquitoes, inhibition of additional miRNAs that do not show differential expression with age will be necessary. We found that the expression levels of a number of the selected predicted targets miR-2946-3p, and miR-317-3p changed due to age. However, further validation of direct interactions of these targets with the corresponding miRNAs needs to be performed.

Previous studies have not found significant changes in the abundances of miRNAs between *Wolbachia w*Mel-infected and uninfected *Drosophila* and mosquito cell lines ^86 87^ or in live flies ^88^. In *Ae. aegypti* Aag2-*w*Mel cells, RT-qPCR was used to assess the abundances of 29 miRNAs that either had been shown in other studies to change in abundance upon *Wolbachia* infection or had potential targets to bind to the 5’ or 3’ ends of the DENV genome ^86^. No statistically significant changes were observed in the abundances of those miRNAs. In *D. melanogaster* JW18 cells, also infected with *w*Mel strain, no significant alteration in miRNAs was observed compared to control JW18Free cells analysed by small RNA-Seq ^87^. In *D. melanogaster* flies infected with *w*Mel strain, no change was found in the abundances of 17 highly abundant miRNAs assessed by RT-qPCR ^88^. In contrast, our previous study on *Ae. aegypti* mosquitoes infected with *w*MelPop ^84^ demonstrated substantial changes in the abundances of a number of miRNAs in mosquitoes infected with *Wolbachia.* In the current study, *Ae. Aegypti* mosquitos did show modulation of miRNAs in response to *w*AlbB, however the effect was weak in comparison to that previously observed in response to *w*MelPop. The reason that substantial modulation of miRNAs occurs in response *w*MelPop, while limited or no modulation of miRNAs occurs in response to *w*AlbB and *w*Mel could be related to pathogenicity of *w*MelPop and lack of pathogenicity of *w*AlbB and *w*Mel.

The *w*MelPop-CLA strain of *Wolbachia* has been shown to regulate the expression of host and *Wolbachia* genes via the production of small RNAs ^74^. Furthermore, sequence homology of small RNA loci exists among a variety of Supergroup-A *Wolbachia* strains, suggesting conservation within certain *Wolbachia* lineages ^74^. In the current study, a subset of the small RNA-Seq reads within our libraries were found to originate from *w*AlbB, and two loci with high coverage were predicted to form stem-loop structures, similar to those of canonical miRNAs. It is possible that stem-loop structures within *w*AlbB non-coding RNAs could be targeted by the host RNAi pathway. Although preliminary, these findings offer an intriguing line of inquiry for future research.

A limitation of this study was the use of whole mosquitoes, rather than individual tissues. Consequently, tissue-specific miRNAs would be underrepresented in the data, as more widely-expressed miRNAs will have greatly outnumbered them at the whole-insect level. With some miRNAs having very low normalised read counts, it is plausible that investigating certain tissues rather than whole insects may allow investigation of some potentially informative miRNAs that were underrepresented in our data.

Overall, our results show that *w*AlbB infection has a modest effect on the miRNA expression profile of *Ae. aegypti*. Some of the differentially expressed miRNAs reported here have been functionally described in *Drosophila* or mosquitoes as having roles in innate immunity, ageing, reproduction, development, or have been shown to be modulated by pathogens including CHIKV and *w*MelPop. Further, we identified two miRNAs, miR-2946-3p, and miR-317-3p as being potentially involved in longevity in *w*AlbB-infected mosquitoes. The results presented here serve to improve our understanding of the molecular interactions governing the relationship between *Wolbachia* and its transinfected host. Future research in this area should be directed at identifying the gene targets of *w*AlbB-modulated miRNAs, followed by experimental manipulation of miRNAs and target genes to determine the effect on *Ae. aegypti* physiology.

## Acknowledgements

Sassan Asgari is supported by the Australian Research Council (DP190102048). Cameron Bishop is supported by a University of Queensland Research Higher Degree scholarship.

## Supplementary Material legends

**Figure S1: Read-length distribution of small RNA-Seq libraries.**

**Figure S2: Bacterial community composition is consistent between WB2 and WB2.tet mosquitoes**. Results of Kraken analysis of reads that did not map to either **A)** *Ae. aegypti* or *w*AlbB in WB2 mosquitoes, or **B)** *Ae. aegypti* in WB2.tet mosquitoes.

**Figure S3: Per-base small RNA-SEQ read depth of the predicted hairpin corresponding to Peak1.**

**Figure S4: Per-base small RNA-SEQ read depth of the predicted hairpin corresponding to Peak 9.**

**Figure S5: *w*AlbB-derived miRNA-like small RNAs. (A)** Predicted hairpin structures corresponding to two regions of the *w*AlbB NZ_CP031221.1 genome that had high coverage of small RNA-Seq reads in libraries derived from WB2 mosquitos. Genomic coordinates indicate the start and end positions of the putative hairpin structure, identified by two peaks of 18-24nt long small RNA-Seq reads. **(B)** Sequence alignment between each predicted stem-loop structure and *W*snRNA-46 and *W*snRNA-59 from Mayoral et al., 2014. Dots represent homologous positions.

**Supplementary Table S1.** Primers and oligos used in this study.

**Supplementary Table S2:** Small RNA-Seq QC mapping statistics.

**Supplementary Table S3:** Genomic coordinates of precursor and mature miRNA loci in the AaeL5.0 reference genome. The base mean is the mean of counts of all samples, normalized for sequencing depth.

**Supplementary Table S4.** Validation of small RNA-Seq using RT-qPCR for nine miRNAs using 5s ribosomal RNA as reference. Ct, cycle threshold. FC, fold-change. CPM, counts per million.

**Supplementary Table S5.** Peaks in coverage of small RNA-Seq reads aligned to the *w*AlbB NZ_CP031221.1. ‘w2’, ‘w6’, ‘w12’ refer to WB2 2 dpe, WB2 6 dpe, and WB2 12 dpe, respectively. Two intergenic regions containing peaks 1 and 9 showed a pattern of read-coverage reminiscent of precursor miRNA loci, and were predicted to form stem-loop structures (Figs. S3-5A).

