## Supplementary Table 1 for "Analysis of *Aedes aegypti* microRNAs in response to *Wolbachia w*AlbB infection and their potential role in mosquito longevity"

| **Primer/oligo** | **sequence** |
|---|---|
| wsp qF | ATCTTTTATAGCTGGTGGTGGT |
| wsp qR | GGAGTGATAGGCATATCTTCAAT |
| Rps17 qF | CACTCCGAGGTCCGTGGTAT |
| Rps17 qR | GGACACTTCGGGCACGTAGT |
| *Ae. aegypti* 5s ribosomal RNA | CGCGTCAGAATGTGAACT |
| AAEL010793 qF | CCTACGACTTCGATTGGGAACA |
| AAEL010793 qR | ATCGTATGCGATCGTACGCC |
| AAEL006171 qF | CTGGCGTAGTTGAAAAGATG |
| AAEL006171 qR | ATGTCAGCAAACCTTGGCCC |
| AAEL010508 qF | ACGAAAATCGCACGGATGGA |
| AAEL010508 qR | CGTTGGGCTTGTAAGTGGTG |
| AAEL006095 qF | CGGCGATTCGTACATCGTCT |
| AAEL006095 qR | CAGCTGGACGCTGAGGATAG |
| AAEL006113 qF | GGGCATCGATCTTTGGGGAT |
| AAEL006113 qR | CGTAGTCAGCGTGGTTGTCT |
| AAEL003402 qF | TATTGTGCGACGCCAAGAGC |
| AAEL003402 qR | GCCCGAAAGGGCATAGGATA |
| miRNA-190-5p inhibitor | AGAUAUGUUUGAUAUUCUUGGUU |
| miRNA-276b-5p inhibitor | AGCGAGGUAUAGAGUUCCUAU |
| Negative control inhibitor | CAGUACUUUUGUGUAGUACAA |
