## Supplementary Table 2 for "Analysis of *Aedes aegypti* microRNAs in response to *Wolbachia w*AlbB infection and their potential role in mosquito longevity"

| Sample | Raw reads | Passed QC | 18-24nt | Mapped x1 | reads mpd >1 | total mapped | % mapped (18-24nt) |
|---|---|---|---|---|---|---|---|
| SRR12893564 | 13,927,840 | 13,926,548 | 3,970,511 | 2,094,977 | 1,736,845 | 3,831,822 | 96.51% |
| SRR12893579 | 15,457,915 | 15,441,355 | 5,405,159 | 2,582,994 | 2,631,028 | 5,214,022 | 96.46% |
| SRR12893578 | 15,519,324 | 15,514,368 | 3,552,681 | 1,774,756 | 1,641,041 | 3,415,797 | 96.15% |
| SRR12893577 | 15,264,515 | 15,258,858 | 4,473,178 | 2,240,998 | 2,017,260 | 4,258,258 | 95.20% |
| SRR12893576 | 14,370,561 | 14,368,661 | 5,347,725 | 2,748,098 | 2,372,449 | 5,120,547 | 95.75% |
| SRR12893575 | 16,493,955 | 16,488,519 | 5,805,852 | 3,004,578 | 2,552,044 | 5,556,622 | 95.71% |
| SRR12893574 | 17,180,689 | 17,167,215 | 5,671,283 | 3,336,000 | 2,119,813 | 5,455,813 | 96.20% |
| SRR12893573 | 10,499,400 | 10,487,552 | 4,817,700 | 2,745,585 | 1,899,599 | 4,645,184 | 96.42% |
| SRR12893572 | 14,454,900 | 14,445,830 | 4,135,898 | 2,341,992 | 1,623,159 | 3,965,151 | 95.87% |
| SRR12893581 | 13,975,176 | 13,945,588 | 5,515,965 | 2,930,216 | 2,192,051 | 5,122,267 | 92.86% |
| SRR12893580 | 12,853,978 | 12,835,587 | 5,248,114 | 2,786,821 | 2,093,335 | 4,880,156 | 92.99% |
| SRR12893571 | 14,962,000 | 14,952,181 | 6,130,774 | 3,179,142 | 2,556,910 | 5,736,052 | 93.56% |
| SRR12893570 | 17,981,052 | 17,970,746 | 6,950,626 | 3,665,646 | 2,845,681 | 6,511,327 | 93.68% |
| SRR12893569 | 16,037,719 | 16,025,114 | 4,623,920 | 2,396,939 | 1,870,612 | 4,267,551 | 92.29% |
| SRR12893568 | 16,675,896 | 16,662,294 | 7,250,301 | 3,706,494 | 2,946,139 | 6,652,633 | 91.76% |
| SRR12893567 | 11,859,587 | 11,833,863 | 4,962,521 | 2,649,644 | 1,906,861 | 4,556,505 | 91.82% |
| SRR12893566 | 15,539,404 | 15,526,359 | 6,272,518 | 3,506,929 | 2,353,557 | 5,860,486 | 93.43% |
| SRR12893565 | 12,698,221 | 12,670,765 | 4,664,429 | 2,464,867 | 1,818,082 | 4,282,949 | 91.82% |
