## Supplementary Table 3 for "Analysis of *Aedes aegypti* microRNAs in response to *Wolbachia w*AlbB infection and their potential role in mosquito longevity"

| miRNA | structure | Chromosome | Start | End | Base mean |
|---|---|---|---|---|---|
| aae-miR-33 | precursor | NC_035107.1 | 50,888,874 | 50,888,954 | -- |
| aae-miR-33-3p | mature | NC_035107.1 | 50,888,881 | 50,888,902 | 46 |
| aae-miR-33-5p | mature | NC_035107.1 | 50,888,926 | 50,888,946 | 28 |
| aae-miR-2940 | precursor | NC_035107.1 | 53,172,117 | 53,172,226 | -- |
| aae-miR-2940-5p | mature | NC_035107.1 | 53,172,121 | 53,172,143 | 14,190 |
| aae-miR-2940-3p | mature | NC_035107.1 | 53,172,204 | 53,172,225 | 28,004 |
| aae-miR-210 | precursor | NC_035107.1 | 54,892,609 | 54,892,677 | -- |
| aae-miR-210-5p | mature | NC_035107.1 | 54,892,618 | 54,892,639 | 172 |
| aae-miR-210-3p | mature | NC_035107.1 | 54,892,653 | 54,892,673 | 4,851 |
| aae-miR-87 | precursor | NC_035107.1 | 80,682,381 | 80,682,488 | -- |
| aae-miR-87-3p | mature | NC_035107.1 | 80,682,402 | 80,682,423 | 3,914 |
| aae-miR-87-5p | mature | NC_035107.1 | 80,682,450 | 80,682,471 | 38 |
| aae-miR-N014 | precursor | NC_035107.1 | 100,807,223 | 100,807,328 | -- |
| aae-miR-N014-3p | mature | NC_035107.1 | 100,807,234 | 100,807,255 | 252 |
| aae-miR-N014-5p | mature | NC_035107.1 | 100,807,296 | 100,807,318 | -- |
| aae-miR-927 | precursor | NC_035107.1 | 115,022,922 | 115,022,998 | -- |
| aae-miR-927-5p | mature | NC_035107.1 | 115,022,930 | 115,022,951 | 5,358 |
| aae-miR-927-3p | mature | NC_035107.1 | 115,022,972 | 115,022,993 | 180 |
| aae-miR-981 | precursor | NC_035107.1 | 118,556,333 | 118,556,424 | -- |
| aae-miR-981-5p | mature | NC_035107.1 | 118,556,349 | 118,556,370 | 16 |
| aae-miR-981-3p | mature | NC_035107.1 | 118,556,391 | 118,556,412 | 6,540 |
| aae-miR-N001 | precursor | NC_035107.1 | 144,289,656 | 144,289,738 | -- |
| aae-miR-N001-5p | mature | NC_035107.1 | 144,289,669 | 144,289,689 | 302 |
| aae-miR-970 | precursor | NC_035107.1 | 157,238,399 | 157,238,507 | -- |
| aae-miR-970-5p | mature | NC_035107.1 | 157,238,424 | 157,238,446 | -- |
| aae-miR-970-3p | mature | NC_035107.1 | 157,238,464 | 157,238,484 | 2,236 |
| aae-miR-252 | precursor | NC_035107.1 | 175,818,151 | 175,818,222 | -- |
| aae-miR-252-3p | mature | NC_035107.1 | 175,818,158 | 175,818,179 | 144 |
| aae-miR-252-5p | mature | NC_035107.1 | 175,818,191 | 175,818,212 | 7,206 |
| aae-miR-929 | precursor | NC_035107.1 | 198,495,510 | 198,495,598 | -- |
| aae-miR-929-5p | mature | NC_035107.1 | 198,495,527 | 198,495,547 | 68 |
| aae-miR-929-3p | mature | NC_035107.1 | 198,495,565 | 198,495,585 | 7 |
| aae-miR-279 | precursor | NC_035107.1 | 207,007,771 | 207,007,866 | -- |
| aae-miR-279-5p | mature | NC_035107.1 | 207,007,789 | 207,007,814 | 18 |
| aae-miR-279-3p | mature | NC_035107.1 | 207,007,832 | 207,007,851 | 17,987 |
| aae-miR-996 | precursor | NC_035107.1 | 207,012,520 | 207,012,618 | -- |
| aae-miR-996-5p | mature | NC_035107.1 | 207,012,537 | 207,012,555 | -- |
| aae-miR-996-3p | mature | NC_035107.1 | 207,012,584 | 207,012,603 | 297 |
| aae-miR-11900 | precursor | NC_035107.1 | 223,944,057 | 223,944,163 | -- |
| aae-miR-11900-5p | mature | NC_035107.1 | 223,944,071 | 223,944,095 | -- |
| aae-miR-11900-3p | mature | NC_035107.1 | 223,944,129 | 223,944,152 | 1,722 |
| aae-miR-999 | precursor | NC_035107.1 | 252,222,287 | 252,222,366 | -- |
| aae-miR-999-5p | mature | NC_035107.1 | 252,222,294 | 252,222,316 | 20 |
| aae-miR-999-3p | mature | NC_035107.1 | 252,222,333 | 252,222,354 | 51,820 |
| aae-miR-34 | precursor | NC_035107.1 | 302,603,044 | 302,603,173 | -- |
| aae-miR-34-3p | mature | NC_035107.1 | 302,603,058 | 302,603,080 | 839 |
| aae-miR-34-5p | mature | NC_035107.1 | 302,603,138 | 302,603,158 | 105,277 |

| | | | | | |
|---|---|---|---|---|---|
| aae-miR-277 | precursor | NC_035107.1 | 302,603,801 | 302,603,895 | -- |
| aae-miR-277-3p | mature | NC_035107.1 | 302,603,814 | 302,603,835 | 73,536 |
| aae-miR-277-5p | mature | NC_035107.1 | 302,603,863 | 302,603,883 | 152 |
| aae-miR-317 | precursor | NC_035107.1 | 302,667,116 | 302,667,208 | -- |
| aae-miR-317-3p | mature | NC_035107.1 | 302,667,130 | 302,667,154 | 50,047 |
| aae-miR-317-5p | mature | NC_035107.1 | 302,667,171 | 302,667,192 | 1,639 |
| aae-miR-12 | precursor | NC_035107.1 | 305,613,535 | 305,613,657 | -- |
| aae-miR-12-3p | mature | NC_035107.1 | 305,613,549 | 305,613,570 | 616 |
| aae-miR-12-5p | mature | NC_035107.1 | 305,613,618 | 305,613,640 | 2,080 |
| aae-miR-1889 | precursor | NC_035107.1 | 305,613,942 | 305,614,045 | -- |
| aae-miR-1889-3p | mature | NC_035107.1 | 305,613,945 | 305,613,966 | 171 |
| aae-miR-1889-5p | mature | NC_035107.1 | 305,614,020 | 305,614,041 | 6,907 |
| aae-miR-283 | precursor | NC_035107.1 | 305,625,581 | 305,625,684 | -- |
| aae-miR-283-3p | mature | NC_035107.1 | 305,625,601 | 305,625,622 | -- |
| aae-miR-283-5p | mature | NC_035107.1 | 305,625,647 | 305,625,669 | 19,871 |
| aae-miR-1175 | precursor | NC_035107.1 | 307,010,034 | 307,010,118 | -- |
| aae-miR-1175-3p | mature | NC_035107.1 | 307,010,048 | 307,010,068 | 3,686 |
| aae-miR-1175-5p | mature | NC_035107.1 | 307,010,083 | 307,010,106 | 2,190 |
| aae-miR-1174 | precursor | NC_035107.1 | 307,010,195 | 307,010,318 | -- |
| aae-miR-1174-3p | mature | NC_035107.1 | 307,010,201 | 307,010,222 | 21,498 |
| aae-miR-1174-5p | mature | NC_035107.1 | 307,010,287 | 307,010,309 | 150 |
| aae-miR-iab_4 | precursor | NC_035107.1 | 308,965,416 | 308,965,490 | -- |
| aae-miR-iab_4-3p | mature | NC_035107.1 | 308,965,424 | 308,965,447 | -- |
| aae-miR-iab_4-5p | mature | NC_035107.1 | 308,965,460 | 308,965,481 | 354 |
| aae-miR-10 | precursor | NC_035107.1 | 310,530,461 | 310,530,550 | -- |
| aae-miR-10-5p | mature | NC_035107.1 | 310,530,476 | 310,530,497 | 28,567 |
| aae-miR-10-3p | mature | NC_035107.1 | 310,530,516 | 310,530,538 | 3,985 |
| aae-miR-993 | precursor | NC_035107.1 | 310,672,009 | 310,672,110 | -- |
| aae-miR-993-3p | mature | NC_035107.1 | 310,672,022 | 310,672,045 | 114 |
| aae-miR-993-5p | mature | NC_035107.1 | 310,672,072 | 310,672,093 | 186 |
| aae-miR-219 | precursor | NC_035108.1 | 3,715,251 | 3,715,337 | -- |
| aae-miR-219-5p | mature | NC_035108.1 | 3,715,262 | 3,715,284 | 20 |
| aae-miR-219-3p | mature | NC_035108.1 | 3,715,307 | 3,715,329 | 50 |
| aae-miR-N013 | precursor | NC_035108.1 | 30,841,052 | 30,841,132 | -- |
| aae-miR-N013-3p | mature | NC_035108.1 | 30,841,063 | 30,841,084 | -- |
| aae-miR-N013-5p | mature | NC_035108.1 | 30,841,100 | 30,841,122 | -- |
| aae-miR-11897a | precursor | NC_035108.1 | 46,313,953 | 46,314,028 | -- |
| aae-miR-11897a-5p | mature | NC_035108.1 | 46,313,964 | 46,313,985 | -- |
| aae-miR-11897a-3p | mature | NC_035108.1 | 46,313,997 | 46,314,018 | -- |
| aae-miR-190 | precursor | NC_035108.1 | 48,811,855 | 48,811,956 | -- |
| aae-miR-190-3p | mature | NC_035108.1 | 48,811,879 | 48,811,900 | 28 |
| aae-miR-190-5p | mature | NC_035108.1 | 48,811,920 | 48,811,942 | 8,942 |
| aae-miR-11894b | precursor | NC_035108.1 | 58,783,349 | 58,783,409 | -- |
| aae-miR-11894b-3p | mature | NC_035108.1 | 58,783,351 | 58,783,372 | -- |
| aae-miR-11894b-5p | mature | NC_035108.1 | 58,783,388 | 58,783,409 | 9 |
| aae-miR-11894a_5 | precursor | NC_035108.1 | 58,785,157 | 58,785,217 | -- |
| aae-miR-11894a_5-3p | mature | NC_035108.1 | 58,785,159 | 58,785,180 | -- |
| aae-miR-11894a_5-5p | mature | NC_035108.1 | 58,785,196 | 58,785,217 | -- |
| aae-miR-11894a_4 | precursor | NC_035108.1 | 58,785,849 | 58,785,909 | -- |
| aae-miR-11894a_4-3p | mature | NC_035108.1 | 58,785,851 | 58,785,872 | -- |

| | | | | | |
|---|---|---|---|---|---|
| aae-miR-11894a_4-5p | mature | NC_035108.1 | 58,785,888 | 58,785,909 | -- |
| aae-miR-11894a_3 | precursor | NC_035108.1 | 58,786,283 | 58,786,343 | -- |
| aae-miR-11894a_3-3p | mature | NC_035108.1 | 58,786,285 | 58,786,306 | -- |
| aae-miR-11894a_3-5p | mature | NC_035108.1 | 58,786,322 | 58,786,343 | -- |
| aae-miR-11894_2 | precursor | NC_035108.1 | 58,787,425 | 58,787,485 | -- |
| aae-miR-11894a_1 | precursor | NC_035108.1 | 58,787,947 | 58,788,007 | -- |
| aae-miR-11894a_1-3p | mature | NC_035108.1 | 58,787,949 | 58,787,970 | -- |
| aae-miR-11894a_1-5p | mature | NC_035108.1 | 58,787,986 | 58,788,007 | -- |
| aae-miR-276a | precursor | NC_035108.1 | 73,120,439 | 73,120,530 | -- |
| aae-miR-276a-3p | mature | NC_035108.1 | 73,120,453 | 73,120,474 | -- |
| aae-miR-276a-5p | mature | NC_035108.1 | 73,120,495 | 73,120,514 | 209 |
| aae-miR-276b | precursor | NC_035108.1 | 73,509,389 | 73,509,473 | -- |
| aae-miR-276b-3p | mature | NC_035108.1 | 73,509,400 | 73,509,421 | 7 |
| aae-miR-276b-5p | mature | NC_035108.1 | 73,509,443 | 73,509,463 | 114 |
| aae-miR-988 | precursor | NC_035108.1 | 93,351,371 | 93,351,446 | -- |
| aae-miR-988-5p | mature | NC_035108.1 | 93,351,381 | 93,351,402 | 483 |
| aae-miR-988-3p | mature | NC_035108.1 | 93,351,418 | 93,351,439 | 285 |
| aae-miR-281 | precursor | NC_035108.1 | 96,469,343 | 96,469,440 | -- |
| aae-miR-281-3p | mature | NC_035108.1 | 96,469,362 | 96,469,383 | 111,160 |
| aae-miR-281-5p | mature | NC_035108.1 | 96,469,399 | 96,469,420 | 59,517 |
| aae-miR-282 | precursor | NC_035108.1 | 96,911,956 | 96,912,051 | -- |
| aae-miR-282-3p | mature | NC_035108.1 | 96,911,967 | 96,911,988 | -- |
| aae-miR-282-5p | mature | NC_035108.1 | 96,912,019 | 96,912,040 | 26 |
| aae-miR-2b | precursor | NC_035108.1 | 107,235,872 | 107,235,959 | -- |
| aae-miR-2b-3p | mature | NC_035108.1 | 107,235,889 | 107,235,908 | 123 |
| aae-miR-2b-5p | mature | NC_035108.1 | 107,235,924 | 107,235,946 | 136 |
| aae-miR-2a | precursor | NC_035108.1 | 107,237,486 | 107,237,583 | -- |
| aae-miR-2a-3p | mature | NC_035108.1 | 107,237,506 | 107,237,525 | 7,324 |
| aae-miR-2a-5p | mature | NC_035108.1 | 107,237,547 | 107,237,569 | 14 |
| aae-miR-13 | precursor | NC_035108.1 | 107,237,641 | 107,237,725 | -- |
| aae-miR-13-3p | mature | NC_035108.1 | 107,237,650 | 107,237,672 | 5,908 |
| aae-miR-13-5p | mature | NC_035108.1 | 107,237,692 | 107,237,713 | 296 |
| aae-miR-2c | precursor | NC_035108.1 | 107,237,990 | 107,238,068 | -- |
| aae-miR-2c-3p | mature | NC_035108.1 | 107,237,996 | 107,238,017 | 629 |
| aae-miR-2c-5p | mature | NC_035108.1 | 107,238,041 | 107,238,061 | -- |
| aae-miR-71 | precursor | NC_035108.1 | 107,238,282 | 107,238,359 | -- |
| aae-miR-71-3p | mature | NC_035108.1 | 107,238,288 | 107,238,309 | 929 |
| aae-miR-71-5p | mature | NC_035108.1 | 107,238,333 | 107,238,354 | 2,315 |
| aae-miR-9a | precursor | NC_035108.1 | 108,461,937 | 108,462,018 | -- |
| aae-miR-9a-3p | mature | NC_035108.1 | 108,461,988 | 108,462,009 | 95 |
| aae-miR-263b | precursor | NC_035108.1 | 153,371,119 | 153,371,210 | -- |
| aae-miR-263b-3p | mature | NC_035108.1 | 153,371,132 | 153,371,153 | 7 |
| aae-miR-263b-5p | mature | NC_035108.1 | 153,371,174 | 153,371,196 | 4,834 |
| aae-miR-2943a | precursor | NC_035108.1 | 158,468,836 | 158,468,917 | -- |
| aae-miR-2943b | precursor | NC_035108.1 | 158,468,998 | 158,469,069 | -- |
| aae-miR-11899 | precursor | NC_035108.1 | 182,319,033 | 182,319,115 | -- |
| aae-miR-11899-3p | mature | NC_035108.1 | 182,319,042 | 182,319,064 | 22 |
| aae-miR-11899-5p | mature | NC_035108.1 | 182,319,083 | 182,319,104 | 22 |
| aae-miR-137 | precursor | NC_035108.1 | 189,612,044 | 189,612,148 | -- |
| aae-miR-137-5p | mature | NC_035108.1 | 189,612,068 | 189,612,090 | -- |

| | | | | | |
|---|---|---|---|---|---|
| aae-miR-137-3p | mature | NC_035108.1 | 189,612,107 | 189,612,128 | 2,746 |
| aae-miR-7 | precursor | NC_035108.1 | 222,802,529 | 222,802,618 | -- |
| aae-miR-7-5p | mature | NC_035108.1 | 222,802,545 | 222,802,568 | 55,960 |
| aae-miR-7-3p | mature | NC_035108.1 | 222,802,585 | 222,802,606 | 12 |
| aae-miR-932 | precursor | NC_035108.1 | 231,381,139 | 231,381,234 | -- |
| aae-miR-932-5p | mature | NC_035108.1 | 231,381,157 | 231,381,179 | 1,464 |
| aae-miR-932-3p | mature | NC_035108.1 | 231,381,195 | 231,381,216 | 253 |
| aae-miR-11911 | precursor | NC_035108.1 | 256,105,994 | 256,106,072 | -- |
| aae-miR-11911-3p | mature | NC_035108.1 | 256,106,006 | 256,106,027 | 31 |
| aae-miR-11911-5p | mature | NC_035108.1 | 256,106,037 | 256,106,059 | -- |
| aae-miR-11893 | precursor | NC_035108.1 | 257,889,031 | 257,889,109 | -- |
| aae-miR-11893-5p | mature | NC_035108.1 | 257,889,042 | 257,889,062 | 16 |
| aae-miR-11893-3p | mature | NC_035108.1 | 257,889,078 | 257,889,099 | 160 |
| aae-miR-2941-1 | precursor | NC_035108.1 | 268,455,991 | 268,456,097 | -- |
| aae-miR-2941-1-3p | mature | NC_035108.1 | 268,456,011 | 268,456,033 | -- |
| aae-miR-2941-1-5p | mature | NC_035108.1 | 268,456,054 | 268,456,075 | -- |
| aae-miR-2941-2 | precursor | NC_035108.1 | 268,456,300 | 268,456,395 | -- |
| aae-miR-2941-2-3p | mature | NC_035108.1 | 268,456,315 | 268,456,337 | -- |
| aae-miR-2941-2-5p | mature | NC_035108.1 | 268,456,357 | 268,456,376 | -- |
| aae-miR-2946 | precursor | NC_035108.1 | 268,456,429 | 268,456,517 | -- |
| aae-miR-2946-3p | mature | NC_035108.1 | 268,456,442 | 268,456,463 | 2,650 |
| aae-miR-2946-5p | mature | NC_035108.1 | 268,456,481 | 268,456,503 | -- |
| aae-miR-196 | precursor | NC_035108.1 | 289,697,987 | 289,698,051 | -- |
| aae-miR-196-3p | mature | NC_035108.1 | 289,697,989 | 289,698,010 | 18 |
| aae-miR-196-5p | mature | NC_035108.1 | 289,698,030 | 289,698,051 | 258 |
| aae-miR-275 | precursor | NC_035108.1 | 307,135,653 | 307,135,736 | -- |
| aae-miR-275-5p | mature | NC_035108.1 | 307,135,669 | 307,135,691 | 294 |
| aae-miR-275-3p | mature | NC_035108.1 | 307,135,709 | 307,135,730 | 20,849 |
| aae-miR-305 | precursor | NC_035108.1 | 307,145,431 | 307,145,519 | -- |
| aae-miR-305-5p | mature | NC_035108.1 | 307,145,446 | 307,145,469 | 6,582 |
| aae-miR-305-3p | mature | NC_035108.1 | 307,145,484 | 307,145,506 | 182 |
| aae-miR-11895 | precursor | NC_035108.1 | 324,949,484 | 324,949,564 | -- |
| aae-miR-11895-3p | mature | NC_035108.1 | 324,949,495 | 324,949,516 | 23 |
| aae-miR-11895-5p | mature | NC_035108.1 | 324,949,533 | 324,949,554 | 579 |
| aae-miR-133 | precursor | NC_035108.1 | 330,417,047 | 330,417,166 | -- |
| aae-miR-133-5p | mature | NC_035108.1 | 330,417,066 | 330,417,088 | -- |
| aae-miR-133-3p | mature | NC_035108.1 | 330,417,128 | 330,417,149 | 1,927 |
| aae-miR-N007 | precursor | NC_035108.1 | 331,681,388 | 331,681,471 | -- |
| aae-miR-N007-5p | mature | NC_035108.1 | 331,681,399 | 331,681,422 | -- |
| aae-miR-N007-3p | mature | NC_035108.1 | 331,681,438 | 331,681,460 | -- |
| aae-miR-9b | precursor | NC_035108.1 | 335,780,561 | 335,780,653 | -- |
| aae-miR-9b-3p | mature | NC_035108.1 | 335,780,575 | 335,780,596 | 32 |
| aae-miR-9b-5p | mature | NC_035108.1 | 335,780,616 | 335,780,639 | 15,091 |
| aae-miR-79 | precursor | NC_035108.1 | 335,780,865 | 335,780,961 | -- |
| aae-miR-79-3p | mature | NC_035108.1 | 335,780,885 | 335,780,907 | 33 |
| aae-miR-79-5p | mature | NC_035108.1 | 335,780,924 | 335,780,946 | 21 |
| aae-miR-306 | precursor | NC_035108.1 | 335,781,035 | 335,781,164 | -- |
| aae-miR-306-3p | mature | NC_035108.1 | 335,781,050 | 335,781,071 | 23 |
| aae-miR-306-5p | mature | NC_035108.1 | 335,781,125 | 335,781,145 | 9,248 |
| aae-miR-9c | precursor | NC_035108.1 | 335,810,358 | 335,810,441 | -- |

| | | | | | |
|---|---|---|---|---|---|
| aae-miR-9c-3p | mature | NC_035108.1 | 335,810,365 | 335,810,386 | 679 |
| aae-miR-9c-5p | mature | NC_035108.1 | 335,810,415 | 335,810,436 | 13,509 |
| aae-miR-375 | precursor | NC_035108.1 | 337,876,516 | 337,876,632 | -- |
| aae-miR-375-5p | mature | NC_035108.1 | 337,876,533 | 337,876,556 | 25 |
| aae-miR-375-3p | mature | NC_035108.1 | 337,876,594 | 337,876,615 | 1,971 |
| aae-miR-14 | precursor | NC_035108.1 | 344,767,521 | 344,767,694 | -- |
| aae-miR-14-5p | mature | NC_035108.1 | 344,767,573 | 344,767,594 | 53 |
| aae-miR-14-3p | mature | NC_035108.1 | 344,767,616 | 344,767,637 | 111,688 |
| aae-miR-263a | precursor | NC_035108.1 | 348,784,426 | 348,784,523 | -- |
| aae-miR-263a-3p | mature | NC_035108.1 | 348,784,449 | 348,784,470 | 70 |
| aae-miR-263a-5p | mature | NC_035108.1 | 348,784,488 | 348,784,510 | 7,142 |
| aae-miR-1891 | precursor | NC_035108.1 | 360,084,853 | 360,084,940 | -- |
| aae-miR-1891-3p | mature | NC_035108.1 | 360,084,868 | 360,084,889 | 1,816 |
| aae-miR-1891-5p | mature | NC_035108.1 | 360,084,904 | 360,084,925 | 10,135 |
| aae-miR-124 | precursor | NC_035108.1 | 369,307,270 | 369,307,346 | -- |
| aae-miR-124-5p | mature | NC_035108.1 | 369,307,282 | 369,307,302 | 8 |
| aae-miR-124-3p | mature | NC_035108.1 | 369,307,320 | 369,307,339 | 334 |
| aae-miR-2945 | precursor | NC_035108.1 | 374,947,133 | 374,947,215 | -- |
| aae-miR-2945-5p | mature | NC_035108.1 | 374,947,143 | 374,947,166 | 28 |
| aae-miR-2945-3p | mature | NC_035108.1 | 374,947,185 | 374,947,205 | 28,652 |
| aae-miR-100 | precursor | NC_035108.1 | 375,117,321 | 375,117,447 | -- |
| aae-miR-100-5p | mature | NC_035108.1 | 375,117,352 | 375,117,373 | 122,990 |
| aae-miR-100-3p | mature | NC_035108.1 | 375,117,392 | 375,117,413 | 738 |
| aae-miR-let_7 | precursor | NC_035108.1 | 375,131,491 | 375,131,564 | -- |
| aae-miR-let_7-5p | mature | NC_035108.1 | 375,131,498 | 375,131,518 | 29,246 |
| aae-miR-let_7-3p | mature | NC_035108.1 | 375,131,539 | 375,131,560 | -- |
| aae-miR-125 | precursor | NC_035108.1 | 375,131,761 | 375,131,856 | -- |
| aae-miR-125-5p | mature | NC_035108.1 | 375,131,782 | 375,131,803 | 11,130 |
| aae-miR-125-3p | mature | NC_035108.1 | 375,131,820 | 375,131,840 | 116 |
| aae-miR-1 | precursor | NC_035108.1 | 399,501,016 | 399,501,092 | -- |
| aae-miR-1-3p | mature | NC_035108.1 | 399,501,024 | 399,501,045 | 185,601 |
| aae-miR-1-5p | mature | NC_035108.1 | 399,501,063 | 399,501,084 | 57 |
| aae-miR-309b | precursor | NC_035108.1 | 402,152,468 | 402,152,539 | -- |
| aae-miR-309b-3p | mature | NC_035108.1 | 402,152,473 | 402,152,493 | -- |
| aae-miR-2944a | precursor | NC_035108.1 | 402,152,656 | 402,152,718 | -- |
| aae-miR-2944a-3p | mature | NC_035108.1 | 402,152,661 | 402,152,680 | -- |
| aae-miR-2944a-5p | mature | NC_035108.1 | 402,152,693 | 402,152,714 | 14 |
| aae-miR-2944b | precursor | NC_035108.1 | 402,152,793 | 402,152,855 | -- |
| aae-miR-2944b-3p | mature | NC_035108.1 | 402,152,795 | 402,152,818 | 9 |
| aae-miR-2944b-5p | mature | NC_035108.1 | 402,152,831 | 402,152,852 | -- |
| aae-miR-286a | precursor | NC_035108.1 | 402,153,076 | 402,153,173 | -- |
| aae-miR-286a-3p | mature | NC_035108.1 | 402,153,090 | 402,153,111 | -- |
| aae-miR-286a-5p | mature | NC_035108.1 | 402,153,141 | 402,153,161 | -- |
| aae-miR-1890 | precursor | NC_035108.1 | 413,654,953 | 413,655,039 | -- |
| aae-miR-1890-3p | mature | NC_035108.1 | 413,654,962 | 413,654,983 | 1,319 |
| aae-miR-1890-5p | mature | NC_035108.1 | 413,655,002 | 413,655,023 | 19 |
| aae-miR-184 | precursor | NC_035108.1 | 417,599,368 | 417,599,456 | -- |
| aae-miR-184-3p | mature | NC_035108.1 | 417,599,381 | 417,599,402 | 69,644 |
| aae-miR-184-5p | mature | NC_035108.1 | 417,599,420 | 417,599,441 | 33 |
| aae-miR-965 | precursor | NC_035108.1 | 429,778,740 | 429,778,819 | -- |

| | | | | | |
|---|---|---|---|---|---|
| aae-miR-965-5p | mature | NC_035108.1 | 429,778,750 | 429,778,773 | 111 |
| aae-miR-965-3p | mature | NC_035108.1 | 429,778,792 | 429,778,813 | 98 |
| aae-miR-N010 | precursor | NC_035108.1 | 439,707,833 | 439,707,917 | -- |
| aae-miR-N010-3p | mature | NC_035108.1 | 439,707,839 | 439,707,861 | -- |
| aae-miR-N010-5p | mature | NC_035108.1 | 439,707,889 | 439,707,910 | -- |
| aae-miR-11905 | precursor | NC_035108.1 | 459,139,427 | 459,139,496 | -- |
| aae-miR-11905-5p | mature | NC_035108.1 | 459,139,429 | 459,139,450 | -- |
| aae-miR-11905-3p | mature | NC_035108.1 | 459,139,474 | 459,139,496 | -- |
| aae-miR-980 | precursor | NC_035109.1 | 1,054,518 | 1,054,594 | -- |
| aae-miR-980-3p | mature | NC_035109.1 | 1,054,524 | 1,054,545 | 22 |
| aae-miR-980-5p | mature | NC_035109.1 | 1,054,562 | 1,054,585 | 58 |
| aae-miR-315 | precursor | NC_035109.1 | 3,444,752 | 3,444,844 | -- |
| aae-miR-315-5p | mature | NC_035109.1 | 3,444,768 | 3,444,790 | 1,412 |
| aae-miR-315-3p | mature | NC_035109.1 | 3,444,810 | 3,444,831 | -- |
| aae-miR-2796 | precursor | NC_035109.1 | 6,069,674 | 6,069,755 | -- |
| aae-miR-2796-3p | mature | NC_035109.1 | 6,069,686 | 6,069,708 | 6,503 |
| aae-miR-2796-5p | mature | NC_035109.1 | 6,069,724 | 6,069,745 | -- |
| aae-miR-1000 | precursor | NC_035109.1 | 74,204,044 | 74,204,141 | -- |
| aae-miR-1000-3p | mature | NC_035109.1 | 74,204,062 | 74,204,083 | 25 |
| aae-miR-1000-5p | mature | NC_035109.1 | 74,204,099 | 74,204,119 | 471 |
| aae-miR-998 | precursor | NC_035109.1 | 75,489,576 | 75,489,668 | -- |
| aae-miR-998-3p | mature | NC_035109.1 | 75,489,590 | 75,489,610 | 6,521 |
| aae-miR-998-5p | mature | NC_035109.1 | 75,489,632 | 75,489,652 | 79 |
| aae-miR-11 | precursor | NC_035109.1 | 75,489,838 | 75,489,952 | -- |
| aae-miR-11-3p | mature | NC_035109.1 | 75,489,865 | 75,489,886 | 109,070 |
| aae-miR-11-5p | mature | NC_035109.1 | 75,489,905 | 75,489,928 | 1,272 |
| aae-miR-971 | precursor | NC_035109.1 | 82,678,939 | 82,679,027 | -- |
| aae-miR-971-5p | mature | NC_035109.1 | 82,678,954 | 82,678,975 | -- |
| aae-miR-971-3p | mature | NC_035109.1 | 82,678,994 | 82,679,016 | 342 |
| aae-miR-92b | precursor | NC_035109.1 | 93,418,505 | 93,418,588 | -- |
| aae-miR-92b-3p | mature | NC_035109.1 | 93,418,516 | 93,418,537 | 1,124 |
| aae-miR-92b-5p | mature | NC_035109.1 | 93,418,555 | 93,418,577 | 46 |
| aae-miR-92a | precursor | NC_035109.1 | 93,477,587 | 93,477,669 | -- |
| aae-miR-92a-3p | mature | NC_035109.1 | 93,477,597 | 93,477,618 | 684 |
| aae-miR-92a-5p | mature | NC_035109.1 | 93,477,637 | 93,477,658 | 18 |
| aae-miR-2765 | precursor | NC_035109.1 | 101,895,441 | 101,895,528 | -- |
| aae-miR-2765-3p | mature | NC_035109.1 | 101,895,455 | 101,895,477 | -- |
| aae-miR-2765-5p | mature | NC_035109.1 | 101,895,491 | 101,895,512 | 94 |
| aae-miR-31 | precursor | NC_035109.1 | 136,174,140 | 136,174,218 | -- |
| aae-miR-31-5p | mature | NC_035109.1 | 136,174,151 | 136,174,171 | 2,812 |
| aae-miR-31-3p | mature | NC_035109.1 | 136,174,186 | 136,174,206 | 218 |
| aae-miR-bantam | precursor | NC_035109.1 | 178,082,826 | 178,082,897 | -- |
| aae-miR-bantam-5p | mature | NC_035109.1 | 178,082,831 | 178,082,854 | 3,881 |
| aae-miR-bantam-3p | mature | NC_035109.1 | 178,082,873 | 178,082,895 | 118,306 |
| aae-miR-N011 | precursor | NC_035109.1 | 240,317,264 | 240,317,340 | -- |
| aae-miR-N011-5p | mature | NC_035109.1 | 240,317,272 | 240,317,292 | -- |
| aae-miR-N011-3p | mature | NC_035109.1 | 240,317,315 | 240,317,335 | -- |
| aae-miR-278 | precursor | NC_035109.1 | 243,123,879 | 243,123,960 | -- |
| aae-miR-278-3p | mature | NC_035109.1 | 243,123,890 | 243,123,910 | 11,340 |
| aae-miR-278-5p | mature | NC_035109.1 | 243,123,930 | 243,123,952 | 77 |

| | | | | | |
|---|---|---|---|---|---|
| aae-miR-307 | precursor | NC_035109.1 | 244,794,355 | 244,794,446 | -- |
| aae-miR-307-3p | mature | NC_035109.1 | 244,794,413 | 244,794,433 | 62 |
| aae-miR-11898a | precursor | NC_035109.1 | 276,079,731 | 276,079,789 | -- |
| aae-miR-11898a-5p | mature | NC_035109.1 | 276,079,733 | 276,079,753 | -- |
| aae-miR-11898a-3p | mature | NC_035109.1 | 276,079,769 | 276,079,789 | -- |
| aae-miR-309a | precursor | NC_035109.1 | 309,335,445 | 309,335,537 | -- |
| aae-miR-309a-3p | mature | NC_035109.1 | 309,335,453 | 309,335,473 | 44 |
| aae-miR-309a-5p | mature | NC_035109.1 | 309,335,497 | 309,335,520 | -- |
| aae-miR-286b | precursor | NC_035109.1 | 309,335,988 | 309,336,085 | -- |
| aae-miR-286b-3p | mature | NC_035109.1 | 309,335,998 | 309,336,020 | 101 |
| aae-miR-286b-5p | mature | NC_035109.1 | 309,336,056 | 309,336,076 | -- |
| aae-miR-316 | precursor | NC_035109.1 | 311,919,545 | 311,919,636 | -- |
| aae-miR-316-5p | mature | NC_035109.1 | 311,919,564 | 311,919,585 | 2,537 |
| aae-miR-316-3p | mature | NC_035109.1 | 311,919,605 | 311,919,626 | -- |
| aae-miR-10365 | precursor | NC_035109.1 | 320,775,611 | 320,775,704 | -- |
| aae-miR-10365-3p | mature | NC_035109.1 | 320,775,622 | 320,775,643 | 67 |
| aae-miR-10365-5p | mature | NC_035109.1 | 320,775,673 | 320,775,694 | 1,248 |
| aae-miR-8 | precursor | NC_035109.1 | 328,527,946 | 328,528,025 | -- |
| aae-miR-8-5p | mature | NC_035109.1 | 328,527,956 | 328,527,977 | 5,071 |
| aae-miR-8-3p | mature | NC_035109.1 | 328,527,995 | 328,528,017 | 293,522 |
| aae-miR-193 | precursor | NC_035109.1 | 360,501,785 | 360,501,871 | -- |
| aae-miR-193-5p | mature | NC_035109.1 | 360,501,796 | 360,501,817 | 43 |
| aae-miR-193-3p | mature | NC_035109.1 | 360,501,839 | 360,501,860 | 11 |
| aae-miR-989 | precursor | NC_035109.1 | 366,439,055 | 366,439,170 | -- |
| aae-miR-989-3p | mature | NC_035109.1 | 366,439,082 | 366,439,103 | 71,200 |
| aae-miR-989-5p | mature | NC_035109.1 | 366,439,124 | 366,439,147 | 108 |
| aae-miR-11898b | precursor | NC_035109.1 | 381,026,876 | 381,026,934 | -- |
| aae-miR-11898b-5p | mature | NC_035109.1 | 381,026,878 | 381,026,898 | -- |
| aae-miR-11898b-3p | mature | NC_035109.1 | 381,026,914 | 381,026,934 | -- |
| aae-miR-2942 | precursor | NC_035109.1 | 390,116,452 | 390,116,534 | -- |
| aae-miR-2942-3p | mature | NC_035109.1 | 390,116,460 | 390,116,482 | 34 |
| aae-miR-2942-5p | mature | NC_035109.1 | 390,116,497 | 390,116,525 | -- |
| aae-miR-957 | precursor | NC_035109.1 | 398,466,618 | 398,466,696 | -- |
| aae-miR-957-5p | mature | NC_035109.1 | 398,466,625 | 398,466,647 | -- |
| aae-miR-957-3p | mature | NC_035109.1 | 398,466,670 | 398,466,691 | 6,360 |
| aae-miR-308 | precursor | NC_035109.1 | 403,338,480 | 403,338,556 | -- |
| aae-miR-308-5p | mature | NC_035109.1 | 403,338,525 | 403,338,546 | 1,736 |
| aae-miR-285 | precursor | NC_035109.1 | 406,875,020 | 406,875,124 | -- |
| aae-miR-285-3p | mature | NC_035109.1 | 406,875,047 | 406,875,068 | 3,050 |
| aae-miR-285-5p | mature | NC_035109.1 | 406,875,086 | 406,875,108 | 38 |
