## Supplementary Table 4 for "Analysis of *Aedes aegypti* microRNAs in response to *Wolbachia w*AlbB infection and their potential role in mosquito longevity"

| miR | sample | Ct Tet 5s | Ct wAlbB 5s | $Log_2FC$ 5s | CPM Tet RNA-Seq | CPM wAlbB RNA-Seq | $Log_2FC$ RNA-Seq |
|---|---|---|---|---|---|---|---|
| miR-184-3p | 2a | 0.009 | 0.012 | 0.457 | 30086.72 | 21683.43 | -0.473 |
| miR-184-3p | 2b | 0.016 | 0.018 | 0.145 | 23221.31 | 24299.58 | 0.065 |
| miR-184-3p | 2c | 0.010 | 0.015 | 0.630 | 22508.70 | 21626.34 | -0.058 |
| miR-184-3p | 6a | 0.014 | 0.019 | 0.390 | 32567.13 | 19805.09 | -0.718 |
| miR-184-3p | 6b | 0.015 | 0.036 | 1.230 | 33438.72 | 19300.74 | -0.793 |
| miR-184-3p | 6c | 0.007 | 0.016 | 1.192 | 32832.30 | 21851.47 | -0.587 |
| miR-184-3p | 12a | 0.017 | 0.029 | 0.752 | 24933.55 | 34228.03 | 0.457 |
| miR-184-3p | 12b | 0.012 | 0.018 | 0.655 | 33762.11 | 30179.70 | -0.162 |
| miR-184-3p | 12c | 0.016 | 0.032 | 0.944 | 26798.87 | 33585.99 | 0.326 |
| miR-190-5p | 2a | 0.000 | 0.000 | 2.636 | 2509.97 | 3615.57 | 0.527 |
| miR-190-5p | 2b | 0.000 | 0.000 | 1.443 | 3101.17 | 3461.22 | 0.158 |
| miR-190-5p | 2c | 0.000 | 0.000 | 2.134 | 1212.08 | 3742.09 | 1.626 |
| miR-190-5p | 6a | 0.000 | 0.000 | 1.104 | 3096.70 | 4766.74 | 0.622 |
| miR-190-5p | 6b | 0.000 | 0.000 | 2.837 | 1374.58 | 5014.42 | 1.867 |
| miR-190-5p | 6c | 0.000 | 0.000 | 1.134 | 3392.17 | 3950.37 | 0.220 |
| miR-190-5p | 12a | 0.000 | 0.000 | 1.961 | 4055.78 | 3533.31 | -0.199 |
| miR-190-5p | 12b | 0.000 | 0.000 | 0.970 | 3555.78 | 3889.45 | 0.129 |
| miR-190-5p | 12c | 0.000 | 0.000 | 2.704 | 3862.88 | 3848.92 | -0.005 |
| miR-276b-5p | 2a | 0.000 | 0.000 | 1.516 | 40.87 | 72.63 | 0.829 |
| miR-276b-5p | 2b | 0.000 | 0.001 | 1.210 | 58.17 | 85.84 | 0.561 |
| miR-276b-5p | 2c | 0.000 | 0.001 | 2.921 | 37.08 | 82.97 | 1.162 |
| miR-276b-5p | 6a | 0.000 | 0.000 | 2.975 | 63.86 | 87.03 | 0.447 |
| miR-276b-5p | 6b | 0.000 | 0.001 | 2.337 | 53.96 | 74.04 | 0.456 |
| miR-276b-5p | 6c | 0.000 | 0.001 | 1.468 | 77.81 | 97.04 | 0.319 |
| miR-276b-5p | 12a | 0.000 | 0.000 | -0.488 | 103.90 | 111.71 | 0.105 |
| miR-276b-5p | 12b | 0.000 | 0.000 | 1.339 | 95.74 | 98.63 | 0.043 |
| miR-276b-5p | 12c | 0.000 | 0.000 | 1.719 | 100.45 | 124.16 | 0.306 |
| miR-2941-1-3p | 2a | 0.001 | 0.000 | -2.837 | 20.22 | 11.85 | -0.771 |
| miR-2941-1-3p | 2b | 0.003 | 0.000 | -3.179 | 14.02 | 8.95 | -0.647 |
| miR-2941-1-3p | 2c | 0.002 | 0.000 | -2.631 | 13.69 | 9.04 | -0.599 |
| miR-2941-1-3p | 6a | 0.002 | 0.002 | -0.478 | 7.69 | 6.97 | -0.141 |
| miR-2941-1-3p | 6b | 0.003 | 0.002 | -0.945 | 8.04 | 10.64 | 0.405 |
| miR-2941-1-3p | 6c | 0.003 | 0.002 | -0.411 | 6.31 | 10.21 | 0.695 |
| miR-2941-1-3p | 12a | 0.002 | 0.003 | 0.400 | 6.51 | 15.04 | 1.208 |
| miR-2941-1-3p | 12b | 0.002 | 0.003 | 0.464 | 3.96 | 8.73 | 1.142 |
| miR-2941-1-3p | 12c | 0.004 | 0.003 | -0.610 | 3.21 | 11.68 | 1.863 |
| miR-308-5p | 2a | 0.000 | 0.000 | 0.261 | 575.30 | 438.89 | -0.390 |
| miR-308-5p | 2b | 0.001 | 0.001 | 0.153 | 476.21 | 439.99 | -0.114 |
| miR-308-5p | 2c | 0.000 | 0.001 | 0.752 | 585.22 | 397.62 | -0.558 |
| miR-308-5p | 6a | 0.001 | 0.001 | 0.692 | 829.02 | 530.99 | -0.643 |
| miR-308-5p | 6b | 0.001 | 0.001 | 0.616 | 912.30 | 435.70 | -1.066 |
| miR-308-5p | 6c | 0.001 | 0.001 | 0.694 | 790.31 | 510.73 | -0.630 |
| miR-308-5p | 12a | 0.001 | 0.002 | 0.872 | 860.03 | 670.63 | -0.359 |
| miR-308-5p | 12b | 0.001 | 0.001 | 0.490 | 954.22 | 806.46 | -0.243 |
| miR-308-5p | 12c | 0.001 | 0.001 | 0.507 | 986.13 | 853.10 | -0.209 |
| miR-31-3p | 2a | 0.004 | 0.007 | 0.801 | 43.95 | 123.44 | 1.490 |
| miR-31-3p | 2b | 0.004 | 0.013 | 1.557 | 92.16 | 107.76 | 0.226 |

| | | | | | | | |
|---|---|---|---|---|---|---|---|
| miR-31-3p | 2c | 0.003 | 0.012 | 1.872 | 76.43 | 121.31 | 0.666 |
| miR-31-3p | 6a | 0.001 | 0.001 | 0.350 | 57.95 | 80.32 | 0.471 |
| miR-31-3p | 6b | 0.002 | 0.002 | -0.052 | 65.44 | 88.08 | 0.429 |
| miR-31-3p | 6c | 0.001 | 0.001 | 0.047 | 82.44 | 89.51 | 0.119 |
| miR-31-3p | 12a | 0.002 | 0.001 | -0.955 | 82.19 | 69.96 | -0.232 |
| miR-31-3p | 12b | 0.001 | 0.001 | 0.024 | 94.16 | 77.97 | -0.272 |
| miR-31-3p | 12c | 0.002 | 0.001 | -0.370 | 88.52 | 68.35 | -0.373 |
| miR-34-5p | 2a | 0.013 | 0.007 | -0.849 | 27140.77 | 21783.18 | -0.317 |
| miR-34-5p | 2b | 0.023 | 0.009 | -1.323 | 16827.98 | 17866.52 | 0.086 |
| miR-34-5p | 2c | 0.018 | 0.010 | -0.867 | 25961.26 | 16777.92 | -0.630 |
| miR-34-5p | 6a | 0.028 | 0.037 | 0.433 | 44573.74 | 44187.88 | -0.013 |
| miR-34-5p | 6b | 0.036 | 0.033 | -0.097 | 46879.10 | 39614.15 | -0.243 |
| miR-34-5p | 6c | 0.021 | 0.039 | 0.930 | 41125.76 | 45430.29 | 0.144 |
| miR-34-5p | 12a | 0.042 | 0.054 | 0.376 | 55620.93 | 60299.06 | 0.117 |
| miR-34-5p | 12b | 0.027 | 0.033 | 0.274 | 53802.74 | 62360.43 | 0.213 |
| miR-34-5p | 12c | 0.050 | 0.065 | 0.387 | 49636.71 | 61823.21 | 0.317 |
| miR-2940-5p | 2a | 0.013 | 0.018 | 0.436 | 6635.10 | 9981.70 | 0.589 |
| miR-2940-5p | 2b | 0.014 | 0.032 | 1.146 | 5879.61 | 6367.91 | 0.115 |
| miR-2940-5p | 2c | 0.007 | 0.032 | 2.205 | 5552.17 | 7372.72 | 0.409 |
| miR-2940-5p | 6a | 0.005 | 0.029 | 2.447 | 9381.17 | 12281.40 | 0.389 |
| miR-2940-5p | 6b | 0.007 | 0.050 | 2.789 | 10162.76 | 17800.88 | 0.809 |
| miR-2940-5p | 6c | 0.005 | 0.053 | 3.284 | 7853.51 | 12732.24 | 0.697 |
| miR-2940-5p | 12a | 0.020 | 0.007 | -1.498 | 12546.02 | 11703.10 | -0.100 |
| miR-2940-5p | 12b | 0.011 | 0.027 | 1.313 | 11820.16 | 19155.17 | 0.696 |
| miR-2940-5p | 12c | 0.010 | 0.018 | 0.826 | 11481.30 | 13675.19 | 0.252 |
| miR-309a-3p | 2a | 0.000 | 0.000 | -0.909 | 7.47 | 3.74 | -0.998 |
| miR-309a-3p | 2b | 0.000 | 0.000 | -2.181 | 8.41 | 2.16 | -1.960 |
| miR-309a-3p | 2c | 0.000 | 0.000 | 0.084 | 11.41 | 3.01 | -1.921 |
| miR-309a-3p | 6a | 0.000 | 0.000 | 0.646 | 14.78 | 20.66 | 0.483 |
| miR-309a-3p | 6b | 0.000 | 0.000 | 0.770 | 17.22 | 20.85 | 0.276 |
| miR-309a-3p | 6c | 0.000 | 0.000 | -0.643 | 13.46 | 20.43 | 0.602 |
| miR-309a-3p | 12a | 0.000 | 0.000 | 0.134 | 19.23 | 32.35 | 0.750 |
| miR-309a-3p | 12b | 0.000 | 0.000 | 0.372 | 20.97 | 38.40 | 0.873 |
| miR-309a-3p | 12c | 0.000 | 0.001 | 0.842 | 22.02 | 28.55 | 0.375 |
| miR-2946-3p | 2a | 0.001 | 0.000 | -2.927 | 506.30 | 640.88 | 0.340 |
| miR-2946-3p | 2b | 0.016 | 0.001 | -4.772 | 780.72 | 510.69 | -0.612 |
| miR-2946-3p | 2c | 0.003 | 0.001 | -2.273 | 375.89 | 541.12 | 0.526 |
| miR-2946-3p | 6a | 0.006 | 0.008 | 0.557 | 613.78 | 1235.53 | 1.009 |
| miR-2946-3p | 6b | 0.014 | 0.021 | 0.623 | 477.58 | 1286.26 | 1.429 |
| miR-2946-3p | 6c | 0.019 | 0.017 | -0.188 | 990.94 | 1168.23 | 0.237 |
| miR-2946-3p | 12a | 0.016 | 0.030 | 0.846 | 1123.97 | 1613.20 | 0.521 |
| miR-2946-3p | 12b | 0.013 | 0.024 | 0.885 | 1613.31 | 1573.64 | -0.036 |
| miR-2946-3p | 12c | 0.021 | 0.041 | 0.974 | 1336.55 | 1802.25 | 0.431 |
