## Supplementary Table 5 for "Analysis of *Aedes aegypti* microRNAs in response to *Wolbachia w*AlbB infection and their potential role in mosquito longevity"

| Name | timepoint(s) in which peak was detected | Start | End | Length | Gene feature overlapping with peak |
|---|---|---|---|---|---|
| Peak_1 | w2_w6_12 | 75270 | 75320 | 50 | Intergenic |
| Peak_2 | w2 | 366284 | 366294 | 10 | gene-DEJ70_RS01810 |
| Peak_3 | w2_w6_12 | 435829 | 435894 | 65 | gene-DEJ70_RS02130;Name=ssrS |
| Peak_4 | w2 | 501208 | 501226 | 18 | gene-DEJ70_RS02470 |
| Peak_5 | w2_w6 | 547309 | 547328 | 19 | gene-DEJ70_RS02685 |
| Peak_6 | w2_w6_12 | 561122 | 561143 | 21 | gene-DEJ70_RS02745 |
| Peak_7 | w2_w6 | 713377 | 713397 | 20 | gene-DEJ70_RS03405;Name=typA |
| Peak_8 | w2_w6_12 | 752990 | 753012 | 22 | gene-DEJ70_RS03590;product=tRNA-Phe |
| Peak_9 | w2_w6_12 | 783582 | 783604 | 22 | Intergenic |
| Peak_10 | w2_w6_12 | 961047 | 961066 | 19 | gene-DEJ70_RS04590;product=tRNA-Leu |
| Peak_11 | w2 | 963346 | 963361 | 15 | gene-DEJ70_RS04600;pseudogene |
| Peak_12 | w12 | 1008958 | 1008977 | 19 | gene-DEJ70_RS04850 |
| Peak_13 | w2_w6_12 | 1043770 | 1043789 | 19 | gene-DEJ70_RS05085;Name=rplB |
| Peak_14 | w12 | 1068991 | 1069008 | 17 | rna-DEJ70_RS05225;product=5S ribosomal RNA |
| Peak_15 | w2_w6 | 1144140 | 1144159 | 19 | gene-DEJ70_RS05535;Name=putA |
| Peak_16 | w2_w6_12 | 1337501 | 1337520 | 19 | Intergenic |
| Peak_17 | w2_w6_12 | 1367049 | 1367068 | 19 | Intergenic |
| Peak_18 | w2_w6_12 | 1476626 | 1476645 | 19 | gene-DEJ70_RS07100 |
