## Supplementary figures and images for "Analysis of *Aedes aegypti* microRNAs in response to *Wolbachia w*AlbB infection and their potential role in mosquito longevity"

### Supplementary Fig. 1

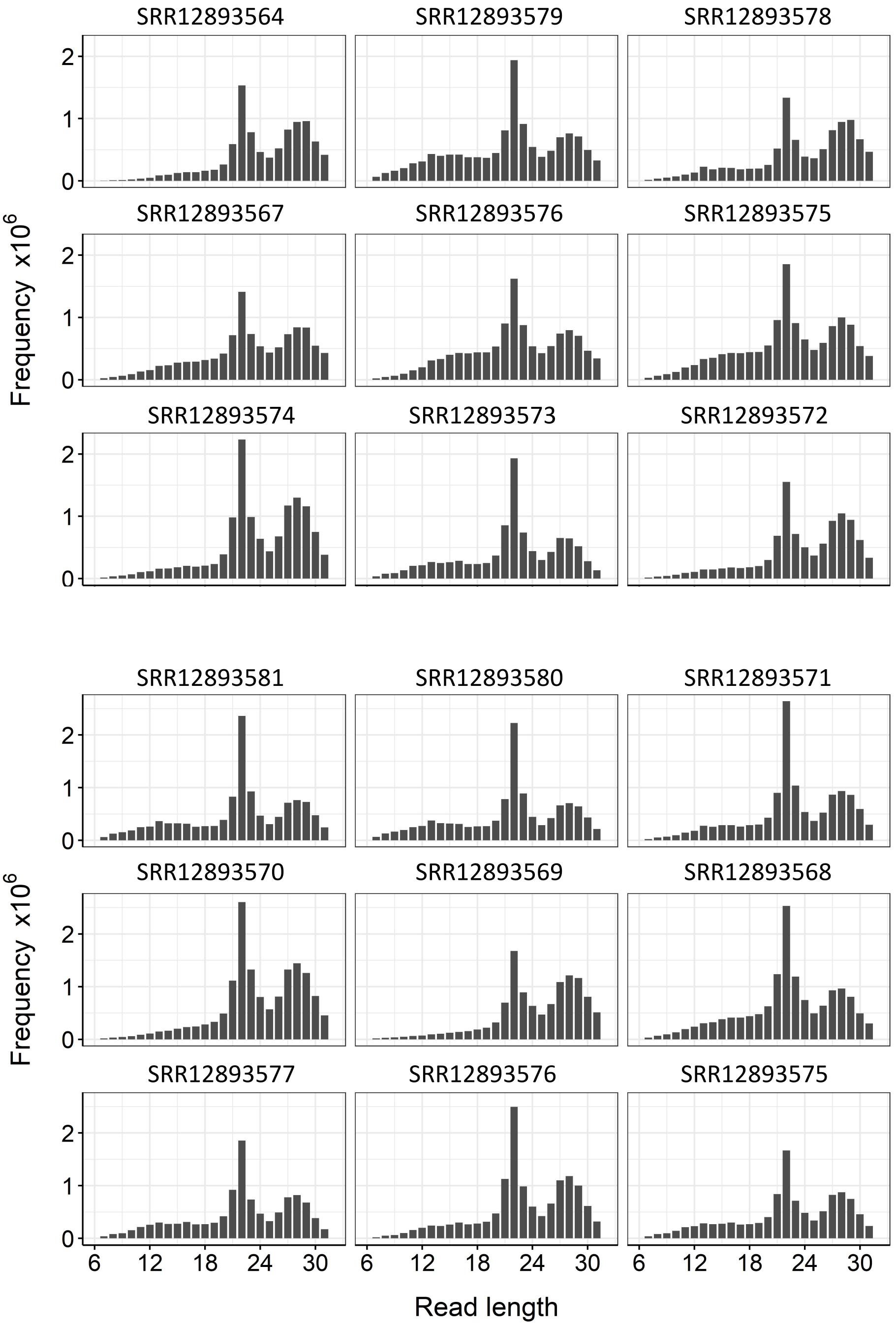

### Supplementary Fig. 2

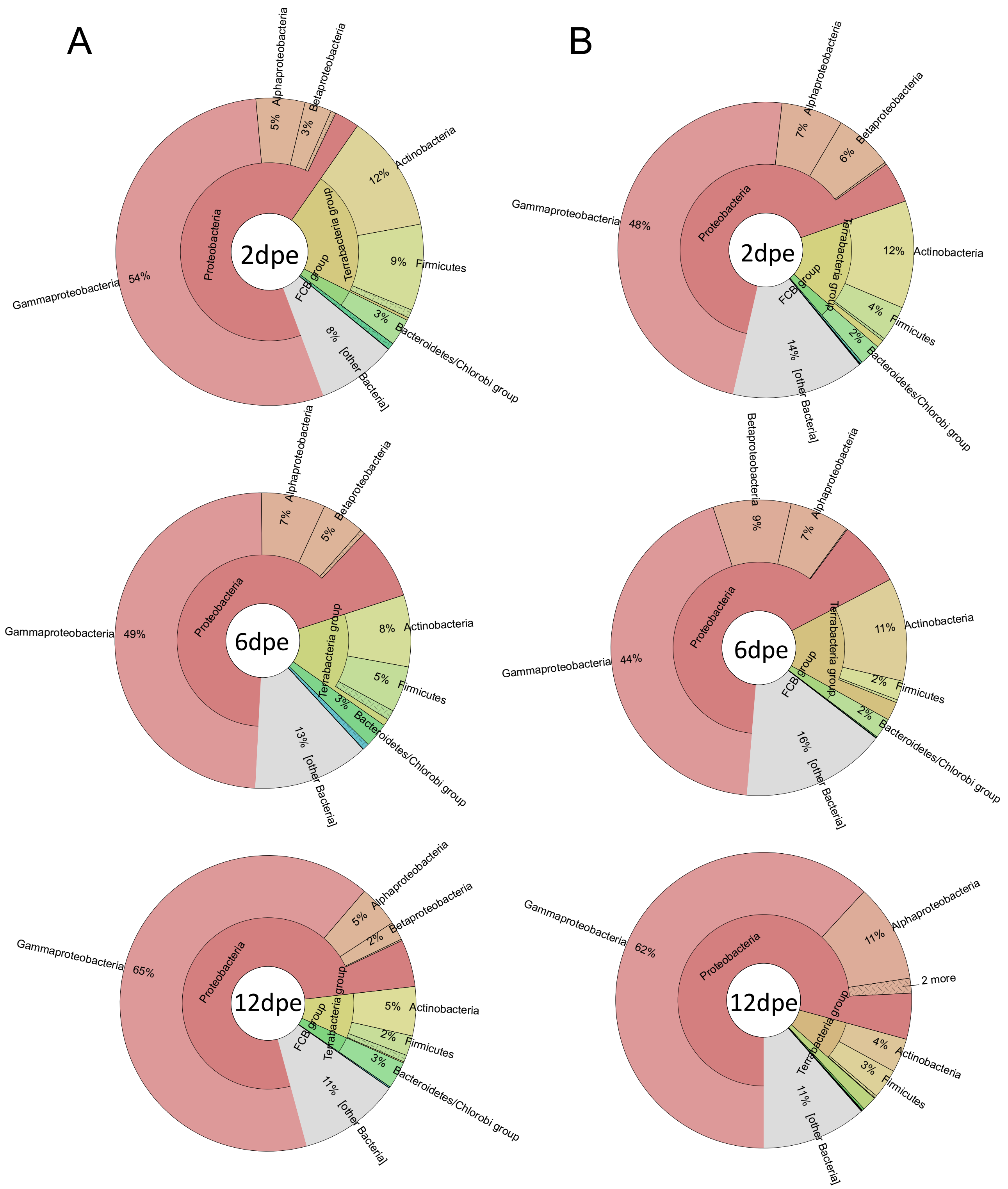

### Supplementary Fig. 3

WB2 2dpi – Peak 1

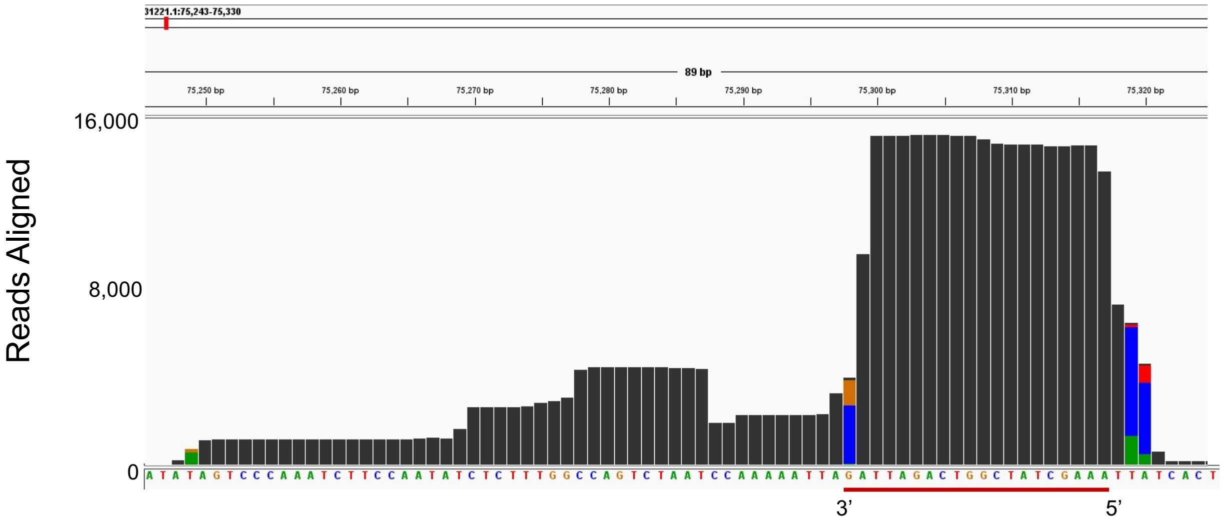


WB2 6dpi – Peak 1

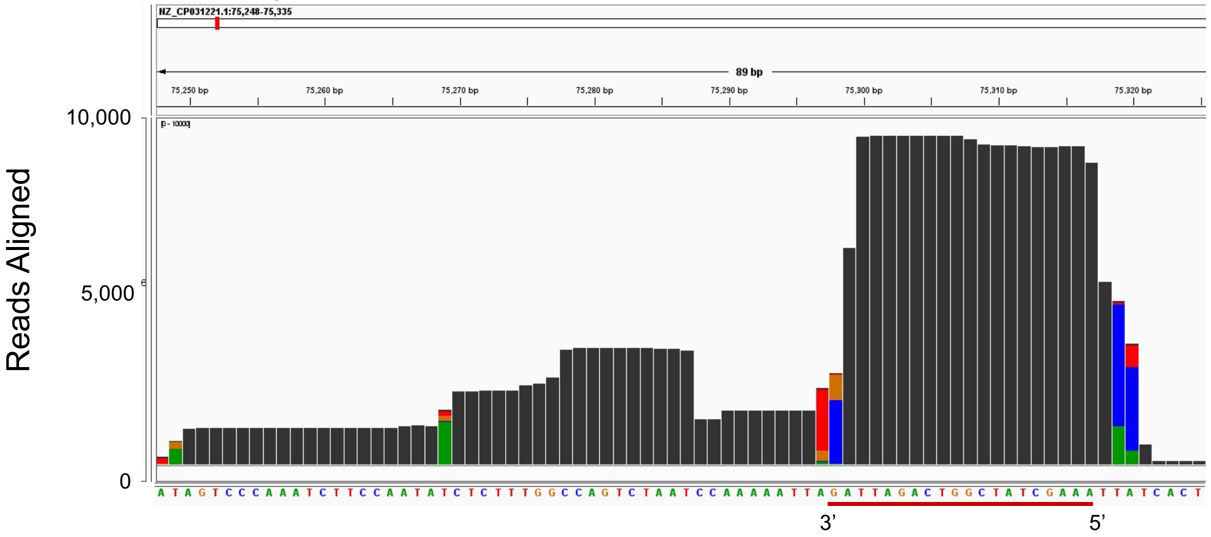


WB2 12dpi – Peak 1

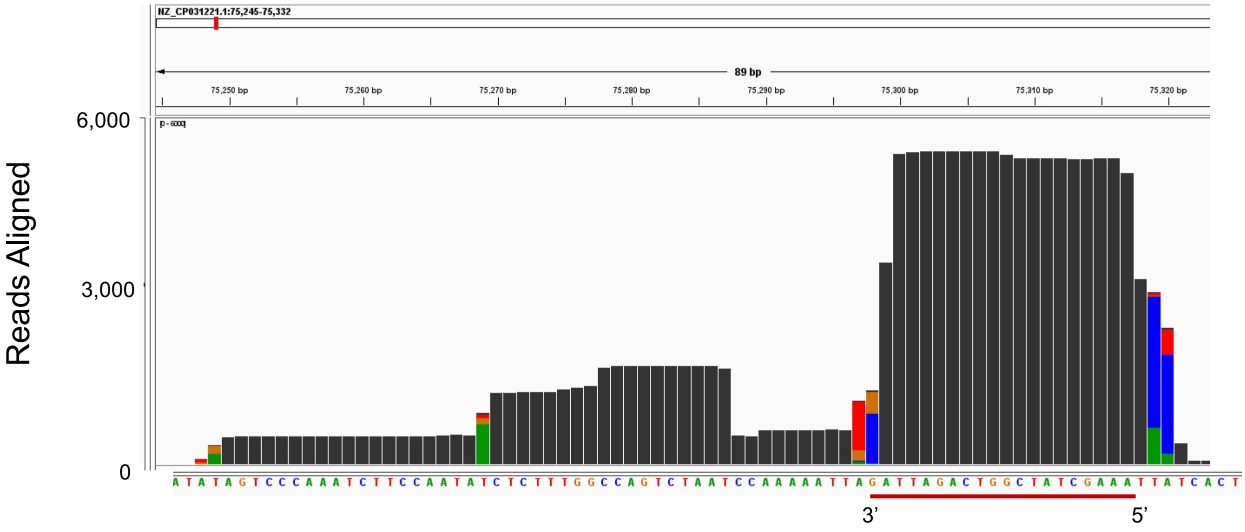

### Supplementary Fig. 4

WB2 2dpi – Peak 9

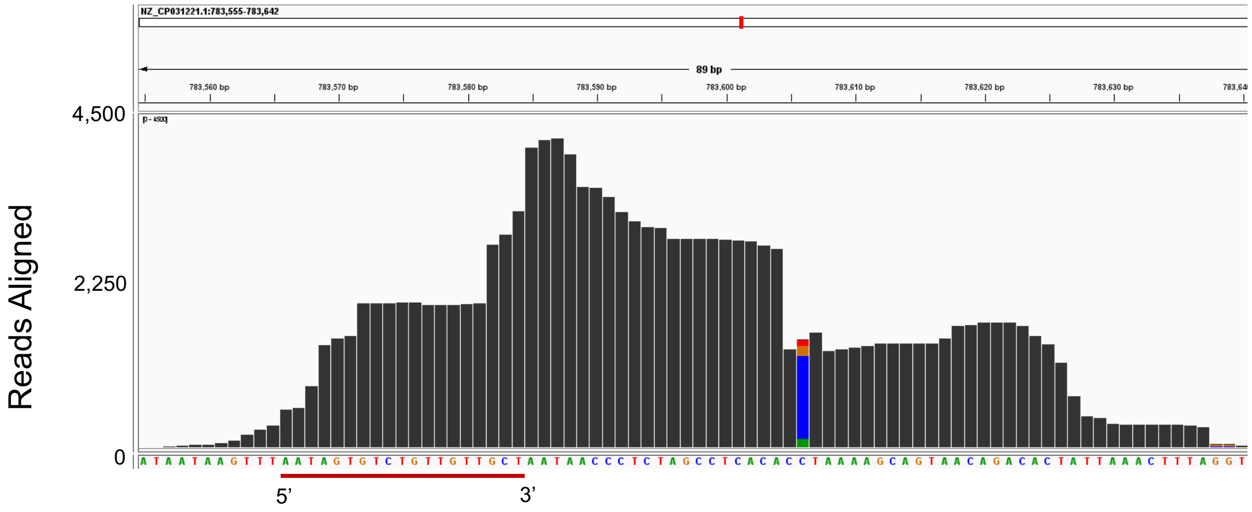


WB2 6dpi – Peak 9

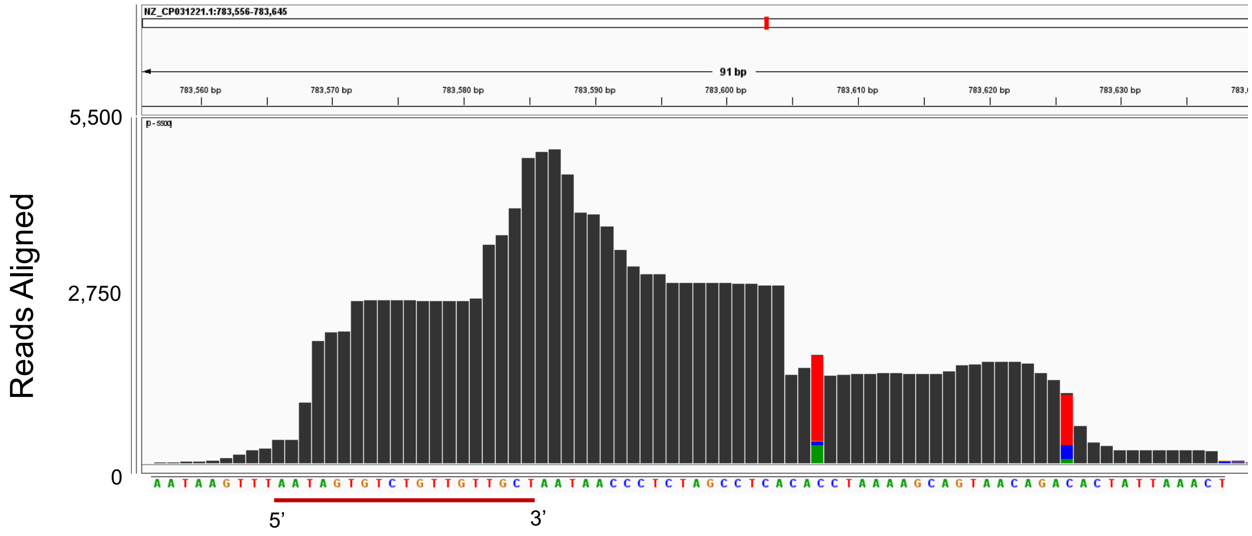


WB2 12dpi – Peak 9

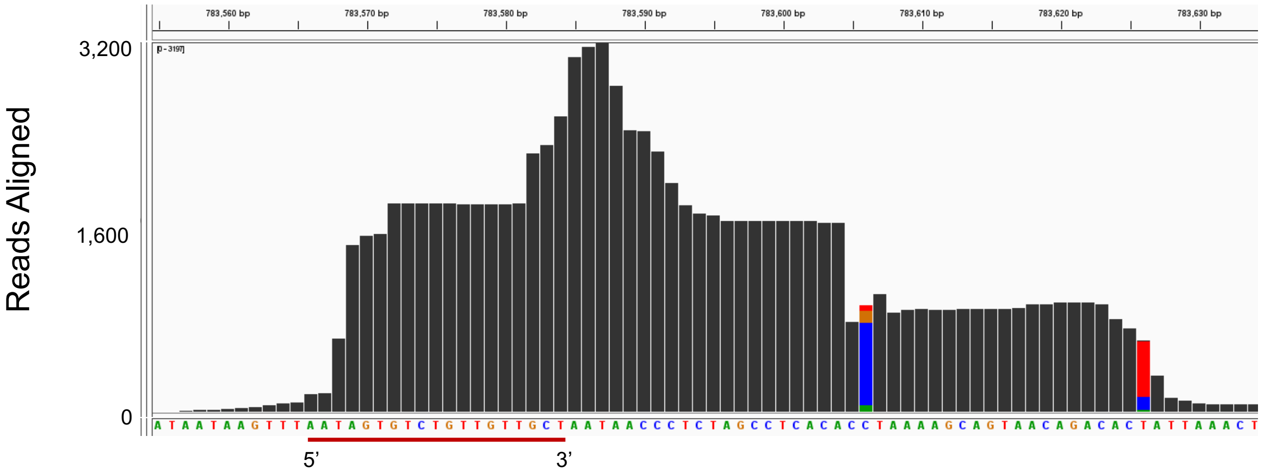
