## Supplementary Fig. 5 for "Analysis of *Aedes aegypti* microRNAs in response to *Wolbachia w*AlbB infection and their potential role in mosquito longevity"

A

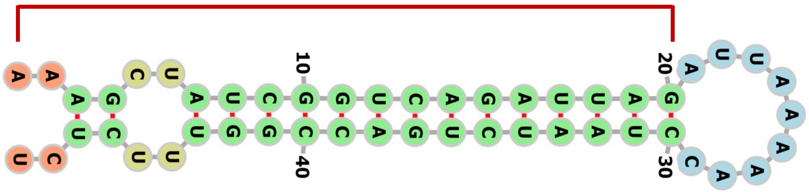

Peak 1 Hairpin structure

75,317 AAAGCTATCGGTCAGATTAGATTAAAAACCTAATCTGACCGGTTTCTCT 75,269

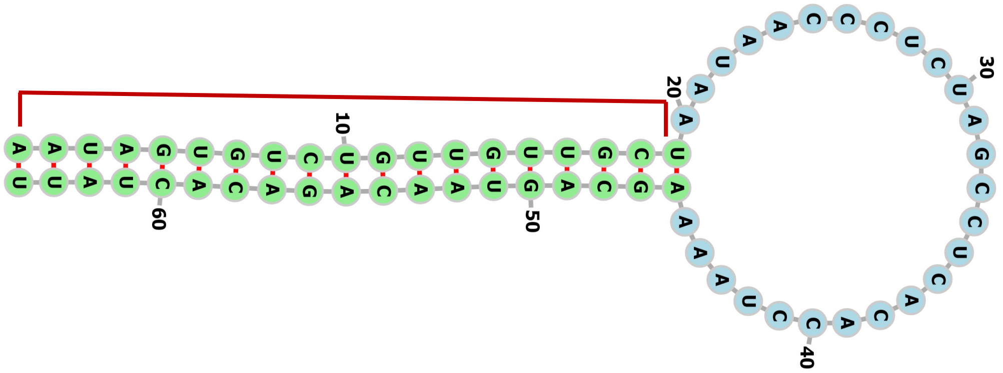

Peak 9 Hairpin structure

783,567 AATAGTGTCTGTTGTTGCTAATAACCCTCTAGCCTCACACCTAAAAGCAG 783,617
783,618 TAACAGACACTATT 783,629

B

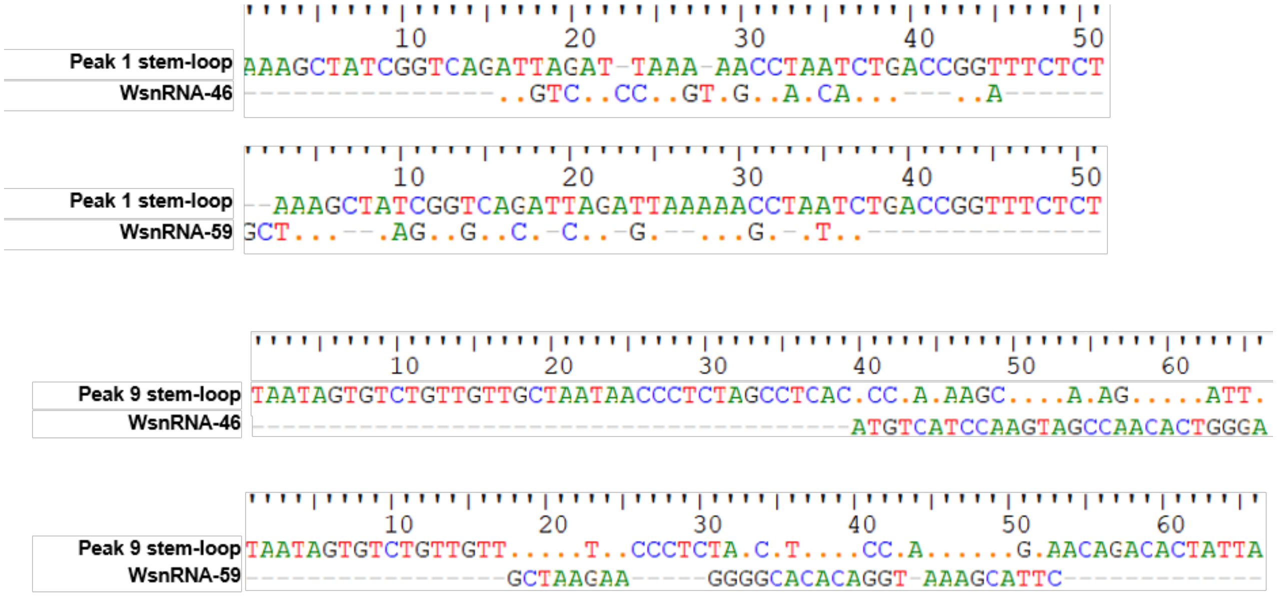
